# Estrogen receptor alpha mutations in breast cancer cells cause gene expression changes through constant activity and secondary effects

**DOI:** 10.1101/2020.04.08.032169

**Authors:** Spencer Arnesen, Zannel Blanchard, Michelle M Williams, Kristofer C Berrett, Zheqi Li, Steffi Oesterreich, Jennifer K Richer, Jason Gertz

## Abstract

While breast cancer patients with tumors that express estrogen receptor α (ER) generally respond well to hormone therapies that block ER activity, a significant number of patients relapse. Approximately 30% of these recurrences harbor activating mutations in the ligand binding domain (LBD) of ER, which have been shown to confer ligand-independent function. However, much is still unclear regarding the effect of mutant ER beyond its estrogen independence. To investigate the molecular effects of mutant ER, we developed multiple isogenic ER mutant cell lines for the most common LBD mutations, Y537S and D538G. These mutations induced differential expression of thousands of genes, the majority of which were mutant allele-specific and were not observed upon estrogen treatment of wildtype cells. These mutant-specific genes showed consistent differential expression across ER mutant lines developed in other laboratories. Wildtype cells with long-term estrogen exposure only exhibited some of these transcriptional changes, suggesting that mutant ER causes novel regulatory effects that are not simply due to constant activity. While ER mutations exhibited minor effects on ER genomic binding, with the exception of ligand independence, ER mutations conferred substantial differences in chromatin accessibility. Mutant ER was bound to approximately a quarter of mutant-enriched accessible regions that were enriched for other DNA binding factors including FOXA1, CTCF, and OCT1. Overall, our findings indicate that mutant ER causes several consistent effects on gene expression, both indirectly and through constant activity.

## Introduction

Estrogen receptor α (ER or ESR1) is a ligand-inducible nuclear hormone receptor that binds estrogens. ER is expressed in roughly 70% of breast cancers and plays a key role in the development and progression of these tumors(1). Due to ER’s role in the growth of ER+ breast cancer, these tumors are generally treated with endocrine therapies, including aromatase inhibitors (AIs), selective estrogen receptor modulators (SERMs), and selective estrogen receptor degraders(SERDs). While hormone therapies have been effective in preventing recurrence, approximately 20% of these cancers develop resistance to hormone therapies and will eventually recur(2,3). Recent genome sequencing efforts have established mutations in the ligand binding domain (LBD) of ER as a common path to hormone therapy resistance that are found in approximately 30% of metastatic ER-positive tumors(4–6).

ER mutations are rarely found in primary ER+ breast cancers but have been observed often in metastatic tumors and in circulating cell free DNA particularly after treatment with AIs(4–10). ER mutations occur almost exclusively in the receptor’s LBD, which is responsible for binding to estrogens and recruiting cofactors, with the majority of mutations occurring at residues Y537 and D538. Crystal structures of the mutant LBD revealed that mutant ER adopts an active conformation in the absence of estrogen binding and several studies have observed ligand-independent transcriptional regulation, growth, and proliferation of ER mutant breast cancer cells both *in vitro* and *in vivo*(4–6,10–16). Large-scale alterations in transcription have also been observed in cells expressing mutant ER(17–19). RNA-seq experiments identified hundreds of ER target genes that are differentially expressed in mutant ER cell lines in the absence of estrogens. A number of genes were also observed as potential “novel” targets of mutant ER. These genes are not regulated by wildtype(WT) ER, but are differentially regulated in mutant ER cells compared to WT regardless of estrogen treatment. Mutant ER also exhibits ligand-independent activity in regard to DNA binding. Recent ChIP-seq experiments showed that mutant ER is capable of binding at many ER binding sites(ERBS) in the absence of estrogens(12,18,20). Additionally, mutant ER may differentially bind at certain sites compared to WT ER(18).

While much has been elucidated regarding the molecular consequences of ER mutations in breast cancer, what drives these effects is less clear. Multiple studies have reported that ER mutations alter the expression of a “novel” or mutant-specific set of genes; however, the consistency of these reported gene expression changes has not been evaluated. Establishing which gene expression changes are found across multiple mutant ER models will assist in identifying molecular or cellular pathways consistently altered when ER is mutated, and reveal potential factors driving mutant ER’s transcriptional impact. In addition, how these mutant-specific transcriptional changes are achieved remains unclear. One model is that mutant-specific gene expression changes are caused by mutant ER’s constant activity. The constitutive activity of mutant ER likely results in the regulation of genes not typically regulated by WT ER with short-term estrogen exposure. These effects could account for many of the mutant-specific expression changes. A second model is that mutations confer neomorphic properties to ER, which could induce new ER genomic binding sites or alter its ability to regulate expression. A third model is that ER mutations cause indirect changes in expression by changing the activity of other transcription factors and/or epigenetic regulators which in turn impact expression. It is unclear which of these models contribute to the novel gene expression observed in breast cancer cells harboring ER mutations.

Here we set out to answer these outstanding questions concerning ER mutations. Using a unique CRISPR/Cas9 strategy that allowed us to introduce the Y537S and D538G mutations and an epitope tag at ER’s endogenous locus, we examined the effects of mutant ER on expression and found thousands of genes consistently impacted by ER mutations across models from different studies. We discovered that approximately half of the mutant ER-specific expression changes can be attributed to constant ER activity. Mutant-specific gene expression changes could be partially explained by alterations in ER genomic binding; however, chromatin accessibility alterations not involving ER were more commonly found nearby mutant-specific genes. Motif analysis of differentially accessible regions identified transcription factors bound at these loci and a role for CTCF and OCT1 in contributing to the unique transcriptional program. Together, our results suggest that ER mutations consistently impact the expression of thousands of novel genes partially through constant ER activity and partially by indirectly altering regulatory elements through additional transcription factors.

## Materials and Methods

### Cell Culture

T-47D cells were obtained from ATCC and MCF-7 cells were obtained from Dr. Jennifer Richer at the University of Colorado Anschutz Medical Campus. WT and mutant T-47D cells were cultured in RPMI 1640 media (Thermo Fisher Scientific) supplemented with 10% fetal bovine serum (Sigma-Aldrich) and 1% penicillin-streptomycin (Thermo Fisher Scientific) (full media). MCF-7 WT and mutant cells were cultured in Minimum Essential Media (MEM, Thermo Fisher Scientific) with 5% fetal bovine serum, 1% penicillin-streptomycin, 1x final concentration non-essential amino acids (NEAA, Thermo Fisher Scientific), and 1nM final concentration insulin (Sigma-Aldrich) (full media). For all experiments, cells were kept at 37°C with 5% CO_2_. Five days before all experiments requiring an E2 induction, cells were placed in hormone-depleted media. Hormone-depleted media consisted of phenol-red free MEM (Thermo Fisher Scientific, 5% charcoal-stripped fetal bovine serum (Thermo Fisher Scientific), 1% penicillin-streptomycin, and NEAA and insulin as described above for MCF-7 cells and phenol-red free RPMI (Thermo Fisher Scientific), 10% charcoal-stripped fetal bovine serum, and 1% penicillin-streptomycin for T-47D cells. Hormone-depleted media was changed twice during each 5-day estrogen-deprivation period in order to ensure complete removal of estrogens from the media. After 5 days of estrogen deprivation, cells were treated with DMSO as a vehicle control (Fisher Scientific) or 10nM E2 (Sigma-Aldrich) for 1 hour for ChIP-seq and ATAC-seq experiments and 8 hours for qPCR and RNA-seq experiments.

### Generation of mutant clones

Mutant clones were generated using the CETCH-seq method described by Savic et al(21). We followed procedures outlined by Blanchard et al.(22) (Supplemental Fig. S1a) using the same guide RNA/Cas9 vector and WT and D538G pFETCH vectors for mutant and WT ER clone generation. For this study, we generated a Y537S pFETCH vector using the same approach as in Blanchard et al., substituting the D538G gBlock with a Y537S gBlock (Supplemental Table S6). Transfections were performed on cells in full media using either Lipofectamin (Thermo Fisher Scientific) or Fugene (Promega) transfection reagents both of which yielded similar success rates. Cells were treated with 1uM SCR7 (Xcessbio) for 3 days post-transfection to block non-homologous end joining. 72 hours post-transfection, cells were treated with G418 (Thermo Fisher Scientific) at 300ug/ml final concentration, which was applied every two days in conjunction with media changes. Single-cell clones were picked and validated using the same procedures as described by Blanchard et al.(22) including limiting dilution plating, colony picking, sanger sequencing and FLAG and ER immunoblots. In all, two clones for each genotype (WT, Y537S, and D538G) were validated for both T-47D and MCF-7 cell lines.

### Proliferation assay

Cells from T-47D WT and mutant clones were cultured in hormone-depleted media for 3 days before plating for proliferation assays. Cells were plated in 96-well plates at approximately 5000 cells per well for each WT or mutant clone. Proliferation was monitored for 48 hours on the IncuCyte Zoom live cell imaging platform (Sartorius) with 10x magnification images obtained at 2 hour intervals. Confluence was measured for three replicates for each cell line in each condition. Linear regression analysis was performed on the log2 confluency percentages that were divided to the initial confluency. Significance differences in proliferation rates were determined using ANOVA to compare the slopes of confluency over time.

### Gene expression analysis

See supplementary materials for experimental and analytical details of quantitative PCR, RNA-seq, siRNA knockdown, and RPPA.

### Chromatin analysis

See supplementary materials for experimental and analytical details of ChIP-seq and ATAC-seq.

### Statistical and Graphical Packages

All statistical analyses were performed in R versions 3.5.2 or 3.5.3(23) except p-values for gene ontology and pathway enrichments which were calculated by Illumina’s BaseSpace Correlation Engine and p-values for motif enrichments which were calculated by MEME suite(24). P-values and statistical tests used can be found throughout the text. Heatmaps for gene expression and differential chromatin accessibility were generated using the pheatmap package in R. Heatmaps display z-scores based on reads per million that align to each gene or region. ChIP-seq results were visualized using the deepTools package(25) to compare constant and mutant-enriched and -depleted ERBS. Distances between mutant up- or down-regulated genes and mutant-enriched or -depleted ERBS or ATAC-seq sites were calculated using a Perl script. Distance plots were generated using the R plot function.

### Data Access

Raw and processed data is available at the Gene Expression Omnibus(GEO) under accession GSE148279.

## Results

### stablishment of endogenously expressed FLAG tagged ER mutant models

Investigation of mutant ER’s molecular and phenotypic consequences relies on the development of models that faithfully recapitulate mutant ER in tumors. Many previous studies investigating mutant ER’s molecular and phenotypic consequences used ectopic expression of mutant ER(4–6,10,14–16,18,26). In order to characterize endogenous ER mutations, we created multiple isogenic clonal lines that heterozygously express FLAG tagged mutant or WT (control) ER from the endogenous locus (22)(Supplemental Fig. S1a). We developed two clones each for WT and the two most common ER LBD mutations (Y537S and D538G) in both T-47D and MCF-7 breast cancer cell lines (Supplemental Fig. S1b). Engineered lines included a FLAG epitope tag at the C-terminus to allow for downstream analyses. The heterozygous expression of FLAG tagged mutant or WT ER and the availability of multiple clones per genotype provided a robust system in which to investigate mutant ER’s molecular effects.

Previous studies have shown estrogen-independent expression of known ER target genes in cells expressing mutant ER(5,12,14,17). To determine whether our mutant clones demonstrated similar gene regulatory effects, we measured the expression of known ER target genes using quantitative PCR in cells treated with 17β-estradiol (E2) or vehicle (DMSO) (Supplemental Fig.S1c). In WT clones, ER regulated genes showed significant changes in expression upon E2 treatment, as expected. In mutant ER clones, these genes were significantly differentially expressed compared to WT controls even with no E2 treatment, while the addition of E2 in some cases further increased the expression of these genes. These results indicate that our mutant clones have ligand-independent function similar to that observed in previous studies(4,5,12,14,17).

### Expression levels of thousands of genes are consistently affected by ER mutations

To investigate whether mutant ER exhibits ligand-independent gene regulation on a genome-wide scale, we performed RNA-seq on WT and mutant clones grown in hormone depleted media and treated with E2 or DMSO for 8 hours. Principal component analysis (PCA) of RNA-seq results indicated that T-47D mutant clones, regardless of E2 treatment, differ distinctly from WT clones in their gene expression profiles (Fig. 1a). Further analysis of T-47D RNA-seq data identified a set of genes that are differentially regulated (adjusted p-value < 0.05) in the WT clones with the addition of E2 (E2-regulated genes). More than half (472 of 771) of the E2-regulated genes in the WT clones were similarly up- or down-regulated in clones from one or both mutants independently of E2 (ligand-independent genes) (Fig. 1b, Supplemental Table S1). A relatively small number of E2-regulated genes were differentially expressed in the opposite direction in ER mutant cells (85 in D538G clones and 32 in Y537S clones). Known E2-regulated genes including *CISH*, *SUSD3*, *KCNK5*, and *PGR* were included in the ligand-independent gene set (example of qPCR validated *CISH* in Fig. 1c). MCF-7 mutants also distinctly clustered apart from MCF-7 WT cells based on RNA-seq data as observed via PCA (Supplemental Fig. S2a). MCF-7 mutant clones exhibited similar ligand-independent effects for both the Y537S and D538G mutations. In these clones, 30% of E2-regulated genes (85 of 289) were regulated in a ligand-independent fashion in one or both mutants (Supplemental Fig. S2b, example of *CISH* in Fig. S2c). These results support previous data showing the ligand-independent function of mutant ER.

**Figure 1.**
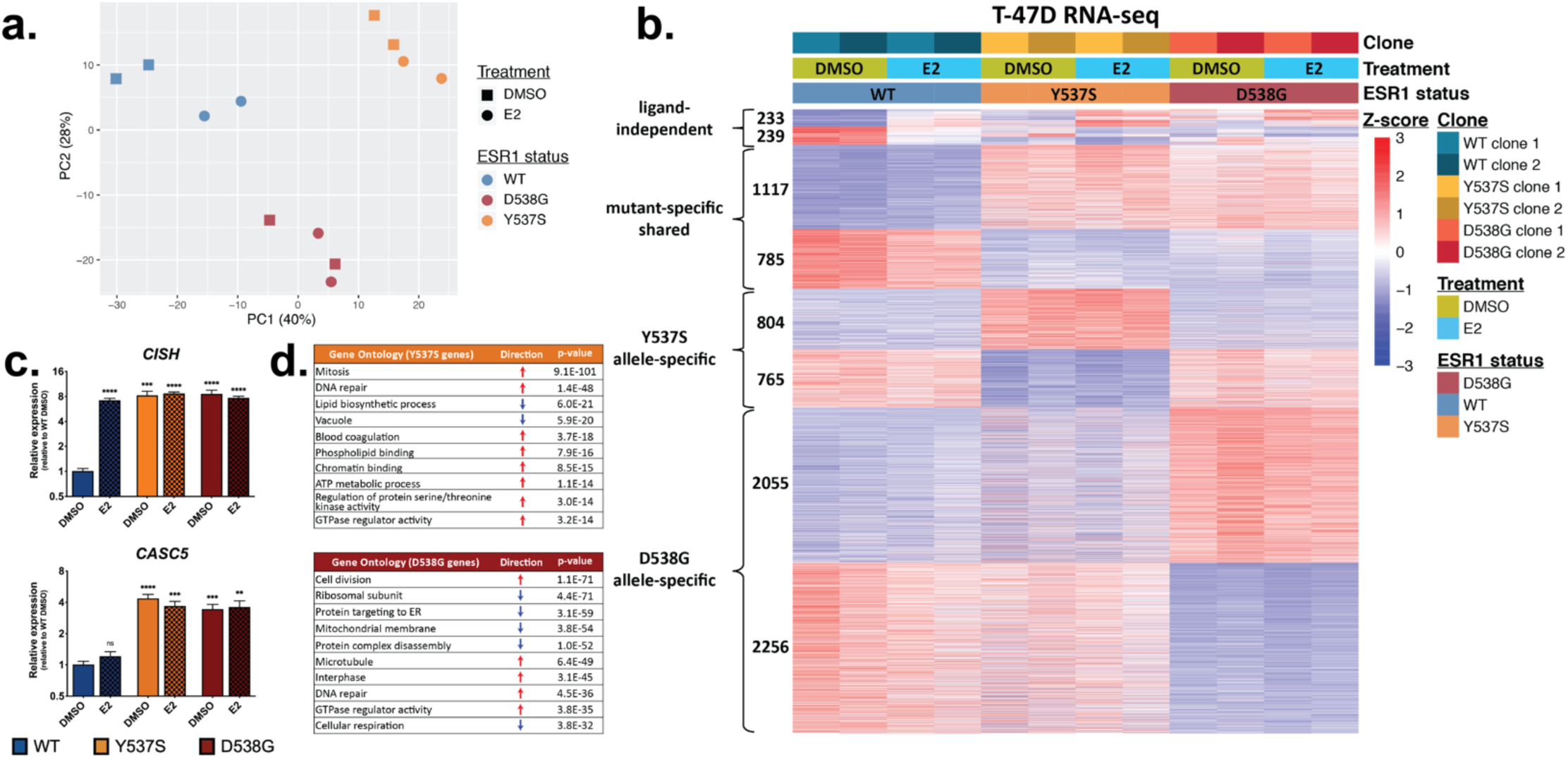
ER mutations exhibit a mutation-specific expression profile. (a) Principal component analysis of RNA-seq data displays the relationships between WT(blue), Y537S(yellow), and D538G(red) T-47D clones. (b) Heatmap shows expression levels for ligand-independent, mutant-specific shared, and allele-specific genes. (c) qPCR validation of ligand-independent (*CISH*) and mutant-specific (*CASC5*) expression. Error bars indicate average ± SEM for two clones for each genotype and each treatment. Student’s two sample t-test was used: ^**^p<0.01, ^***^p<0.001, ^****^p<0.0001, and n.s. = not significant. (d) Enriched gene ontology terms for Y537S- and D538G-specific differentially regulated genes with Fisher’s exact test p-values are shown.

In addition to ligand-independent gene regulation, ER mutant cells also exhibit differential expression of a large number of genes that are not differentially regulated upon an 8-hour E2 treatment in WT ER cells. In T-47D clones, over 1900 genes were up-(1117) or down-regulated (785) in both the Y537S and the D538G mutations (shared mutant-regulated genes) (Fig. 1b, example of qPCR validated *CASC5* in Fig. 1c). The majority of mutant-regulated genes were allele-specific with 1569 Y537S-specific and 4311 D538G-specific up- or down-regulated genes (Fig. 1b). Similar results were seen in our MCF-7 mutant clones, with over a thousand shared mutant-specific genes and thousands of genes regulated in an allele-specific fashion (Supplemental Fig. S2b, examples in Supplemental Fig. S2c). Overall, mutant-specific genes made up the vast majority of differentially regulated genes in our mutant clones, indicating that a major result of ER mutation, in addition to ligand-independent regulation of WT ER target genes, is the differential regulation of a novel gene expression program. Of these mutant-specific genes, nearly 30% of D538G regulated genes (up-regulated genes: p-value = 5.1e^−55^; down-regulated genes: p-value = 1.8e^−59^) and 15% of Y537S differentially regulated genes (up-regulated genes: p-value = 2.3e^−28^; down-regulated genes: p-value = 7.04e^−17^) were shared across both cell lines (T-47D and MCF-7, examples in Supplemental Fig. S2d) suggesting that while some mutant-specific genes are consistently observed in multiple cell lines, many may be cell line-specific.

To investigate enriched pathways in mutation-regulated genes, we performed gene ontology and pathway analysis and found that mutant-specific genes in T-47D cells were highly enriched for positive roles in cell cycle and cell division or protein synthesis and processing (Fig. 1d). Mutant-specific genes in MCF-7 cells were enriched for genes involved in increased cellular migration and motility but decreased cell division (Supplemental Fig. S2e). Using reverse phase protein array (RPPA) analysis, we validated that a number of gene expression changes that occur in our T-47D mutant clones also occur at the protein level (Supplemental Fig. S3a). Many proteins that were significantly different between mutant and WT cells were also differentially regulated at the transcriptional level in our mutant clones according to our RNA-seq results (Supplemental Fig. S3b). These proteins included cyclins and cyclin-dependent kinases as well as DNA-repair proteins which are key to successful cell-cycle progression. In addition to cell cycle-related genes, we also validated increased expression of *IGF1R* at the protein level and decreased expression of *CDH1* which codes for E-cadherin (Supplemental Fig. S3b). These proteins play important roles in cell growth and cell motility, and IGF1R has been shown to be activated by ER mutations (27,28).

To explore phenotypes associated with the observed gene expression changes, we analyzed proliferation in T-47D cells. Using live cell imaging on the IncuCyte Zoom platform, we found that ER mutant cells grow significantly faster in full media than WT lines with the D538G lines exhibiting the fastest growth (Figure S4a). In hormone depleted media, the effects on growth were much more pronounced with all mutant clones growing more rapidly than WT cells (Figure S4b). The increased growth of ER mutant cells may be related to E2F factors, which have been shown to mediate secondary effects of estrogens in breast cancer cells(29,30). E2F factors *E2F1*, *E2F2*, *E2F7*, *E2F8*, and E2F dimerizing partner *TFDP1* were significantly up-regulated in ER mutant T-47D cells. In addition, motif comparison of T-47D ER mutant up-regulated and down-regulated gene promoters showed strong enrichment for E2F motifs in up-regulated gene promoters (p-values = 1.4e^-19^ and 1.8e^-20^ for Y537S and D538G). These observations suggest that E2F factors may play a role in mediating the gene expression effects of ER mutations. The gene expression changes found in the MCF-7 ER mutant cells suggested a change in migratory potential. Consistent with this observation, previous studies have shown that ER mutant MCF-7 cells migrate faster in a scratch wound assay(14). We also found that our ER mutant MCF-7 cells are more capable of invading matrigel (Williams et al. co-submitted; CAN-20-1200R-A). Together, these findings indicate that the gene expression changes observed in ER mutant cells correspond to phenotypic changes. This expanded transcriptional impact may also affect other properties such as anchorage independent growth and metastatic outgrowth as observed in Williams et al. (CAN-20-1200R-A).

We observed increased expression of *ITGA6* (Figure S2d), which is also known as CD49f and is used as a marker of luminal progenitor cells(31,32). This led us to question whether ER mutant cells resemble luminal progenitor cells more than wildtype cells. We compared our mutant-specific up- and down-regulated genes to genes found to be up- or down-regulated in luminal progenitor cells in a previous study(32). For our T-47D mutant-specific genes, we observed marginally significant overlaps with luminal progenitor genes. However, we found that our MCF-7 mutant-specific up-regulated genes exhibited highly significant overlaps with luminal progenitor increased genes (Supplemental Table S2). These results suggest that ER mutations may lead to a more stem-like luminal progenitor expression state, at least in some contexts.

Previous studies have described T-47D and MCF-7 ER mutant lines made using CRISPR or lentiviral techniques and have reported similar E2-independent and mutant-specific gene regulation(12,17–19). In an effort to define which genes are consistently regulated by ER mutations, we compared RNA-seq data from the clones described above with RNA-seq data collected from ER mutant lines described in two other studies(17,18). A PCA plot of these multi-lab data from MCF-7 lines clustered them by study than by ER mutational status (Supplemental Fig. S5a). This is likely due to multiple factors including differences in mutation introduction techniques, duration of hormone depletion, E2 induction time, and sequencing library generation protocol. However, within PCA study clusters, clones segregated by mutational status and E2 treatment. Multivariate analysis of these data to account for variation between studies identified genes that were consistently differentially regulated by ER mutations across all datasets. Over 1900 genes were up-(993) or down-regulated (910) by both mutations across all MCF-7 lines (Fig. 2a, examples in Fig. 2b, Supplemental Table S3). Thousands of genes were specific to each mutation (1378 D538G-specific genes and 2678 Y537S-specific genes). Genes that were consistently differentially expressed in mutant lines from all three studies were enriched for a variety of gene ontology terms shared between both mutations including innate immune response, mitochondrial matrix, and extracellular matrix. Allele specific terms were also observed including cellular respiration and ribosomal subunit for the Y537S mutation and cell-cell adhesion and lipid biosynthesis for the D538G mutation (Fig. 2c). Although these terms differ somewhat from those identified in individual studies’ lines, they represent underlying changes that consistently occur in response to the expression of ER mutations and therefore may provide direction for future studies regarding mutant ER action and effects.

**Figure 2.**
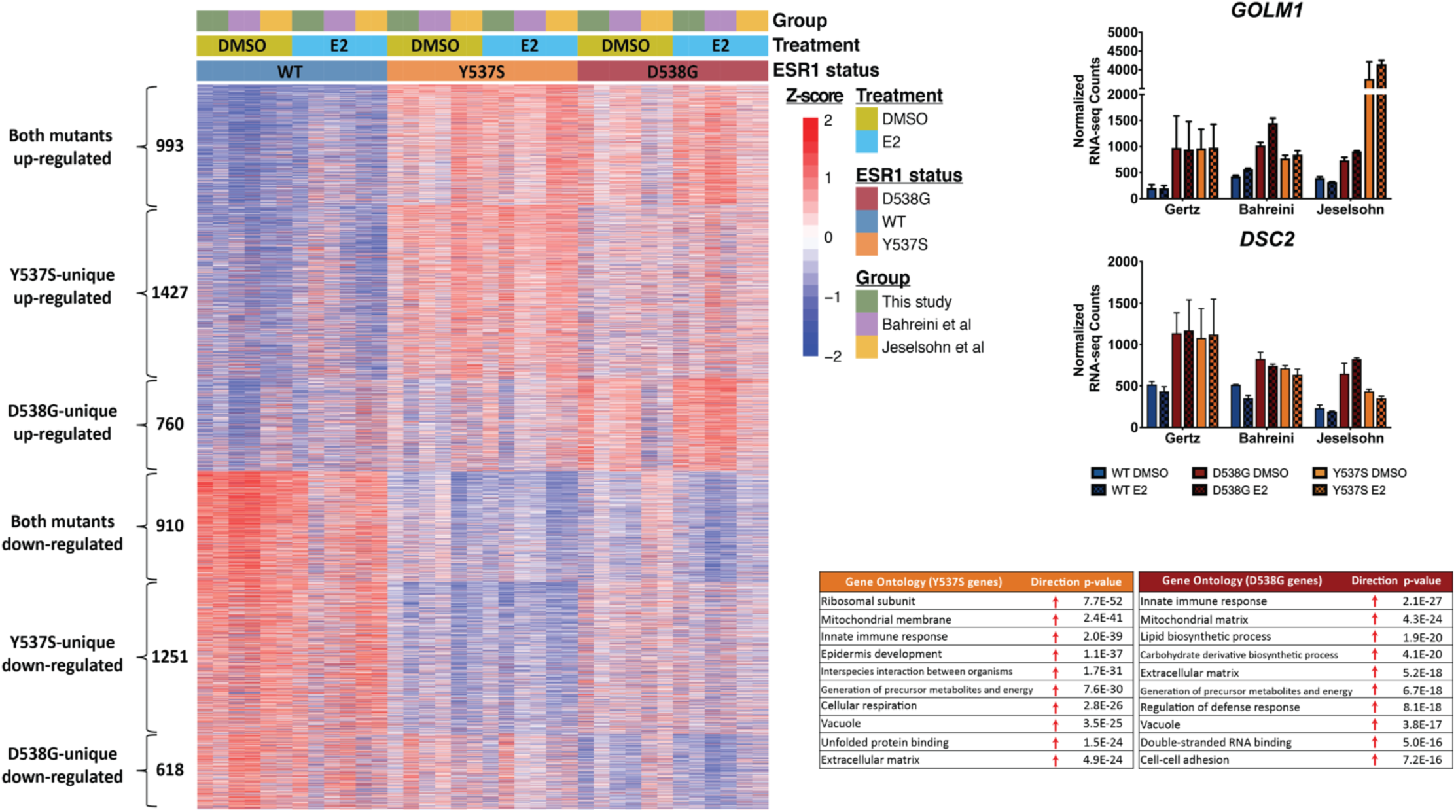
Gene expression analysis from three independent studies reveals consistent ER mutant-specific patterns. (a) Heatmap shows levels of consistent mutant-specific genes. (b) Examples of genes exhibiting consistent mutant-specific gene expression are shown. (c) Significantly enriched gene ontology terms are displayed for Y537S-specific and D538G-specific gene sets from the multi-study comparison.

We used WT and mutant T-47D RNA-seq data from the same three studies to perform an identical multivariate analysis as above with the exception that one study lacked the D538G mutation in T-47D cells. The T-47D multivariate analysis yielded similar results to the MCF-7 analysis with over 1700 up- or down-regulated genes shared between both mutations and thousands of additional genes differentially regulated by each mutant allele uniquely (Supplemental Fig. S5b, examples in Supplemental S5c). These genes were heavily enriched for cell cycle-related genes (Supplemental Fig. S5d). Overall, the comparison of multiple lines from multiple individual groups provides evidence of consistent regulation of novel genes by ER mutations.

To determine if the expression changes that we observed across studies can be seen in patient samples, we analyzed RNA-seq data from the MET500 study(33). By splitting metastatic breast cancer samples by ER mutation status, we identified 70 up-regulated genes in ER mutant tumors and 160 down-regulated genes in ER mutant tumors at a false discovery rate of 5%. Comparing these gene lists to the multi-study genes described above, we observed significant overlaps of 16-22% of up-regulated genes and 10-13% of down-regulated genes (Supplemental Table S4). The overlaps were more significant between up-regulated genes with T-47D Y537S and MCF-7 D538G showing the highest levels of enrichment. These results indicate that genes whose expression is impacted in well controlled cell line experiments are enriched for differential expression in patient tumors based on ER mutation status.

### Constant ER activity can explain some of mutant ER’s impact on gene expression

The constant activity of ER brought about by LBD mutations could explain some of the mutant-specific gene expression changes observed in our RNA-seq analyses. Long-term constitutive ER activity may lead to the differential expression of genes that are in fact regulated by WT ER but do not change expression with short-term E2 treatment. To determine how much of mutant ER’s gene regulatory changes can be attributed to mutant ER’s constitutive activity, we treated T-47D and MCF-7 WT clones with E2 for 25 days. WT ER cells were collected at 5 day intervals and RNA was harvested for RNA-seq analysis. A PCA of RNA-seq data compared WT T-47D cells treated with long-term (5, 10, 15, 20, 25 days) E2 to WT ER short-term (0 and 8hr) E2 treatments and mutant T-47D cells. PC1, which accounted for 50% of the variance between samples, revealed a distinct separation between long-term E2 treated WT ER cells and both WT ER short-term E2 treatments and mutant ER cells. (Fig. 3a). PC2 grouped long-term E2-treatments apart from short-term E2 treatments and instead clustered them closer to Y537S and D538G mutant cells. Together, these observations indicate that while long-term ER activity may account for some of the differences between mutant and WT cells, it may not account for all mutant ER gene expression changes. Further analysis of RNA-seq data revealed that 54% of all mutant-regulated genes were similarly up- or down-regulated by long-term E2 treatment (adjusted p-value < 0.05) (Fig. 3b,c, example of *TBCD* in Fig. 3d). However, a considerable number of mutant ER regulated genes were not differentially regulated with long-term E2 treatment in WT cells, but remained specific to one or both mutations (example of *COPS2* in Fig. 3d). Similar results were observed in MCF-7 WT cells treated with long-term E2 with 38% of all mutant-specific genes being explained by constant ER activity (Supplemental Fig. S6a-d). These results suggest that while the constitutive activity of the mutant receptor may account for approximately half of the gene expression changes observed in mutant ER breast cancer cells, a large percentage of mutant-specific gene expression changes cannot be attributed to constant ER activity. These unexplained mutant-specific gene expression changes may instead result from additional properties of mutant ER which allow it to directly or indirectly alter the expression of these genes.

**Figure 3.**
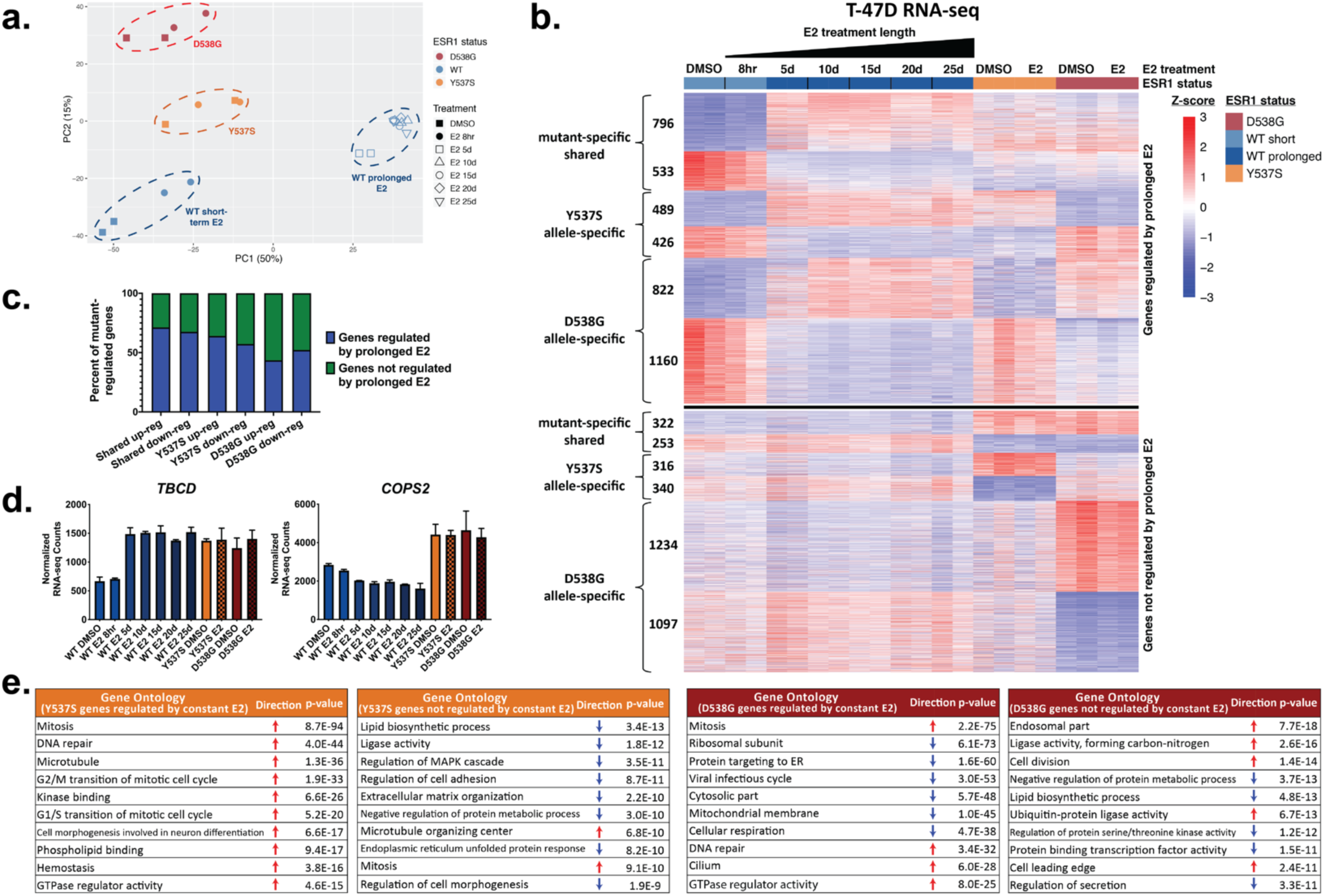
Constant activity of ER explains approximately half of T-47D mutant-specific genes. (a) Principle component analysis of RNA-seq data captures the relationships between T-47D ER mutant clones and WT clones with short- or long-term E2 treatment. (b) Heatmap shows all mutant-specific genes separated by long-term E2-treatment expression (top: genes regulated by long-term E2 treatment; bottom: genes not regulated by long-term E2 treatment). (c) Percent of mutant-specific genes regulated by long-term ER activation in WT clones is shown. (d) Bar graphs show examples of genes that are regulated by long-term E2 treatment (*TBCD*) and that are differentially regulated by mutation only (*COPS2*). (e) Significantly enriched gene ontology terms are displayed for Y537S-specific and D538G-specific gene sets partitioned based on overlap with long-term E2 regulated genes.

### ER mutations do not cause broad reprogramming of genomic binding

Mutant-specific gene expression changes could result from altered genomic binding by mutant ER compared to WT ER. To investigate this possibility, we performed ChIP-seq experiments on WT and mutant T-47D and MCF-7 clones, grown for 5 days in hormone depleted media followed by 1 hour E2 or DMSO treatments, using an antibody that recognizes the FLAG epitope tag. Results from these ChIP-seq experiments compared well with previously performed ER ChIP-seq experiments performed in another lab, with 68-73% binding site overlaps observed between samples of the same *ESR1* genotype(18). In WT T-47D clones, ER binding by WT ER is absent at thousands of genomic regions in the absence of E2 but is gained with the addition of E2 (Fig. 4a, example in Fig. 4b). ChIP-seq of mutant ER showed that for both mutations ER bound to the majority of WT ERBS independent of E2 treatment and mutant ER binding at these sites increased with the addition of E2. The majority of ER bound sites were identified in both mutants and in E2-treated WT clones (34920 constant sites); only ~10% (3837 sites) of ER-bound regions exhibited significant differential binding between WT and mutant with the large majority of these sites being differentially bound in Y537S mutant clones (Fig. 4c, example in Fig. 4d). In both constant and mutant-enriched and -depleted sites the ER binding motif (ERE) was the most highly enriched motif.

**Figure 4.**
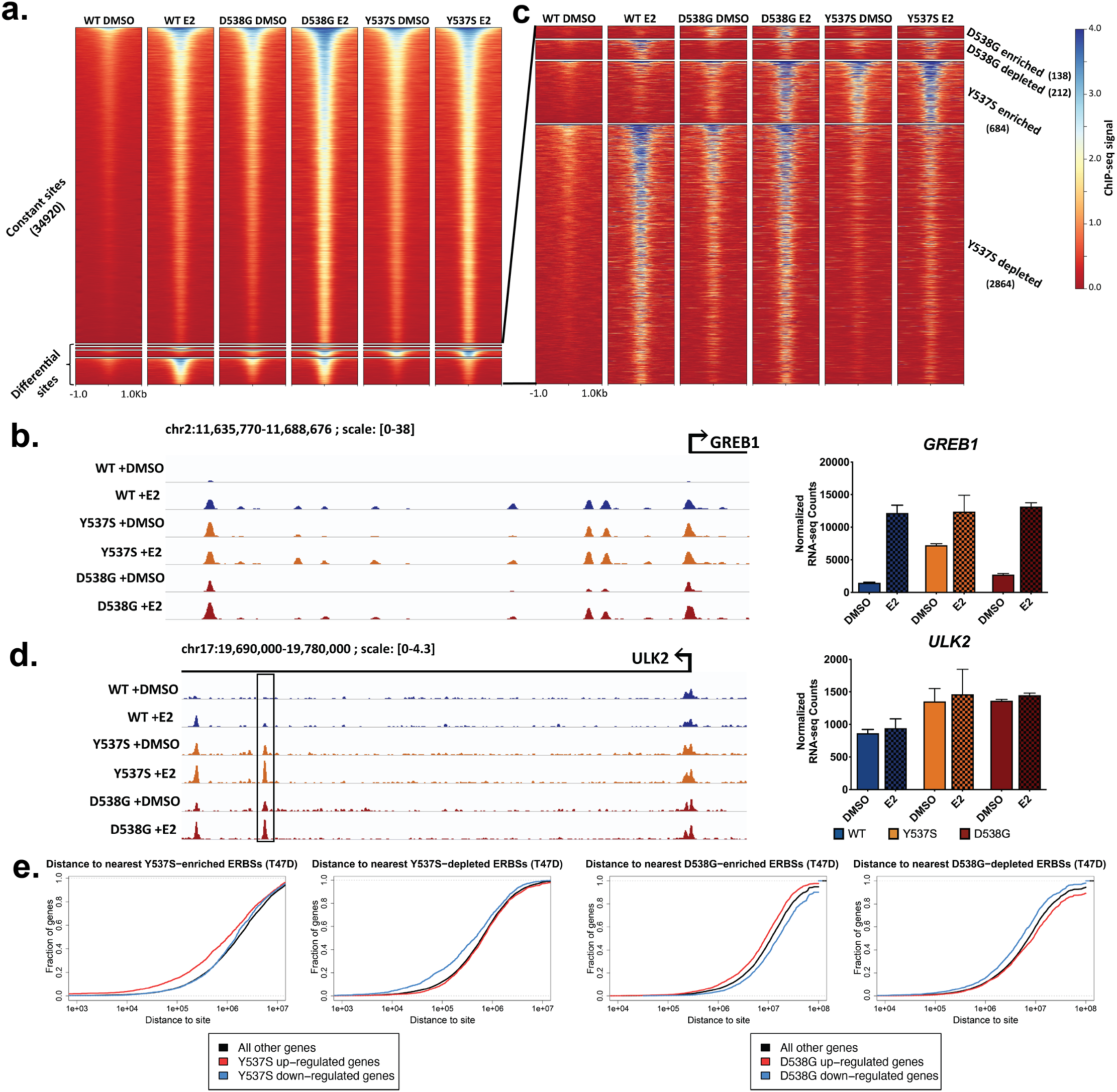
Limited changes in ER’s genomic binding are observed in ER mutant cells. (a) Heatmap shows the binding signal of constant and mutant-specific ERBS in T-47D WT and mutant clones with addition of DMSO or E2. (b) Example locus of ligand-independent (constant) ER-binding, arrow indicates TSS of *GREB1*. All tracks are normalized to the same scale. *GREB1* ligand-independent gene expression is shown on the right. (c) Heatmap displays the signal of mutant-enriched or depleted sites only. (d) Example locus of mutant-enriched ER-binding. All tracks are normalized to the same scale. *ULK2* gene expression is shown on the right. (e) Distance from mutant-specific genes to mutant-enriched and -depleted ERBS is shown as a cumulative distribution.

To determine how much of the mutant-specific expression differences that we observed could be explained by ER binding, we analyzed the distance between mutant-specific genes and constant, mutant-enriched, or mutant-depleted ERBS. Both Y537S and D538G mutant-specific up- and down-regulated genes are significantly closer to constant ERBS than are all other genes (background). Using a previously described distance threshold of 100 kilobasepairs(kb)(34–36), we determined the number of mutant-specific genes that contained ERBS near their transcription start site (TSS). Of both Y537S and D538G mutant-specific genes, 55-80% contain nearby constant ERBS, which is 13-16% more than background genes. Mutant-specific up- and down-regulated genes are also significantly closer to mutant-enriched and -depleted sites respectively (Fig. 4e, Supplemental Table S5). 10% of Y537S up- and down-regulated genes contain at least one nearby Y537S-enriched or -depleted ERBS (Supplemental Table S5). Enriched and depleted ERBS were found to a much lesser extent in D538G mutant clones; less than 1% of D538G-specific up-regulated genes contain a nearby D538G-enriched ERBS and less than 2% of D538G-specific down-regulated genes contain a nearby D538G-depleted ERBS. These results indicate that while constant ligand-independent ER genomic binding likely affects the expression of a portion of mutant-specific genes, differential ER binding, particularly in D538G mutant cells, may only have a subtle effect on gene expression.

Similar results were seen in MCF-7 WT and mutant clones where 36606 sites were bound in WT cells treated with E2 that were also bound in mutant cells regardless of E2 treatment (Supplemental Fig. S7a-c). Differential ER binding in MCF-7 clones also follows similar patterns to those observed in T-47D clones with fewer D538G differentially bound ERBS (1161) than Y537S differentially bound ERBS (3020). Similar numbers of mutant-specific genes harboring an ERBS within 100 kb of the TSS were observed in MCF-7 cells compared to T-47D cells (Supplemental Fig. S7d, Supplemental Table S5). Approximately 13% of Y537S-enriched and -depleted genes contain at least one nearby enriched or depleted Y537S ERBS while only 1-6% of D538G genes contain nearby D538G-enriched or -depleted ERBS. While ligand-independent genomic binding by mutant ER is apparent, the low percentage of ERBS that exhibited significant differential binding suggests that changes in ER binding may not have a broad impact on the regulation of mutant-specific genes. However, mutant-specific genes are significantly closer to mutant-altered ERBS and therefore may account for a small portion (2-14%) of mutant-specific differential gene expression.

### ER mutations cause widespread changes in chromatin accessibility

Because *ESR1* mutations do not appear to drastically reprogram ER genomic binding, we hypothesized that additional changes might occur at genomic loci bound by ER, or other transcription factors, that contribute to mutant ER’s unique gene expression program. In order to identify genomic regions that could contribute to the observed mutant ER gene expression changes, we analyzed genome-wide chromatin accessibility using ATAC-seq in our WT and mutant T-47D clones with a 1-hour E2 or DMSO treatment. Minimal changes were observed in chromatin accessibility in response to E2 treatment with only 169 regions exhibiting significantly increased accessibility and no regions of decreased accessibility in WT clones. However, in mutant ER cells, thousands of genomic regions exhibit increased (1808 in Y537S clones and 1563 in D538G clones; mutant-enriched sites, Fig. 5a, example in Fig. 5b) or decreased accessibility (2060 in Y537S clones and 1587 in D538G clones; mutant-depleted sites, Fig. 5a) when compared to WT clones. The changes in chromatin accessibility were mutation-allele specific with only 21.6% of mutant-enriched and 23.7% of mutant-depleted regions shared between both mutations. To determine the extent to which ER could be directly involved at regions of altered chromatin accessibility, we overlapped ER bound sites identified from our FLAG tag ChIP-seq experiments with all regions of differential chromatin accessibility. Of the mutant-enriched ATAC-seq regions, only 30-50% overlap with ERBS while only 20-30% of mutant-depleted ATAC-seq regions are bound by ER (Fig. 5d), indicating that many of these chromatin changes are not direct effects of mutant ER. To determine if the results seen in our mutant cells are recapitulated in mutant lines described in previous studies, we performed ATAC-seq on T-47D WT and mutant D538G and Y537S cells first described in Bahreini et al(17). We performed multivariate differential accessibility analysis on the combined ATAC-seq data from these sets of mutant ER models and identified thousands of genomic regions that show consistent differential chromatin accessibility between mutant ER and WT lines (Supplemental Fig. S8a). ER binding again occurs at a minority (20-30%) of the differentially accessible regions from the multivariate analysis (Supplemental Fig. S8b). These findings suggest that considerable changes in chromatin accessibility are consistently seen at thousands of sites in multiple independent ER mutant models, and that they are largely mutation-allele specific and mostly lack ER binding.

**Figure 5.**
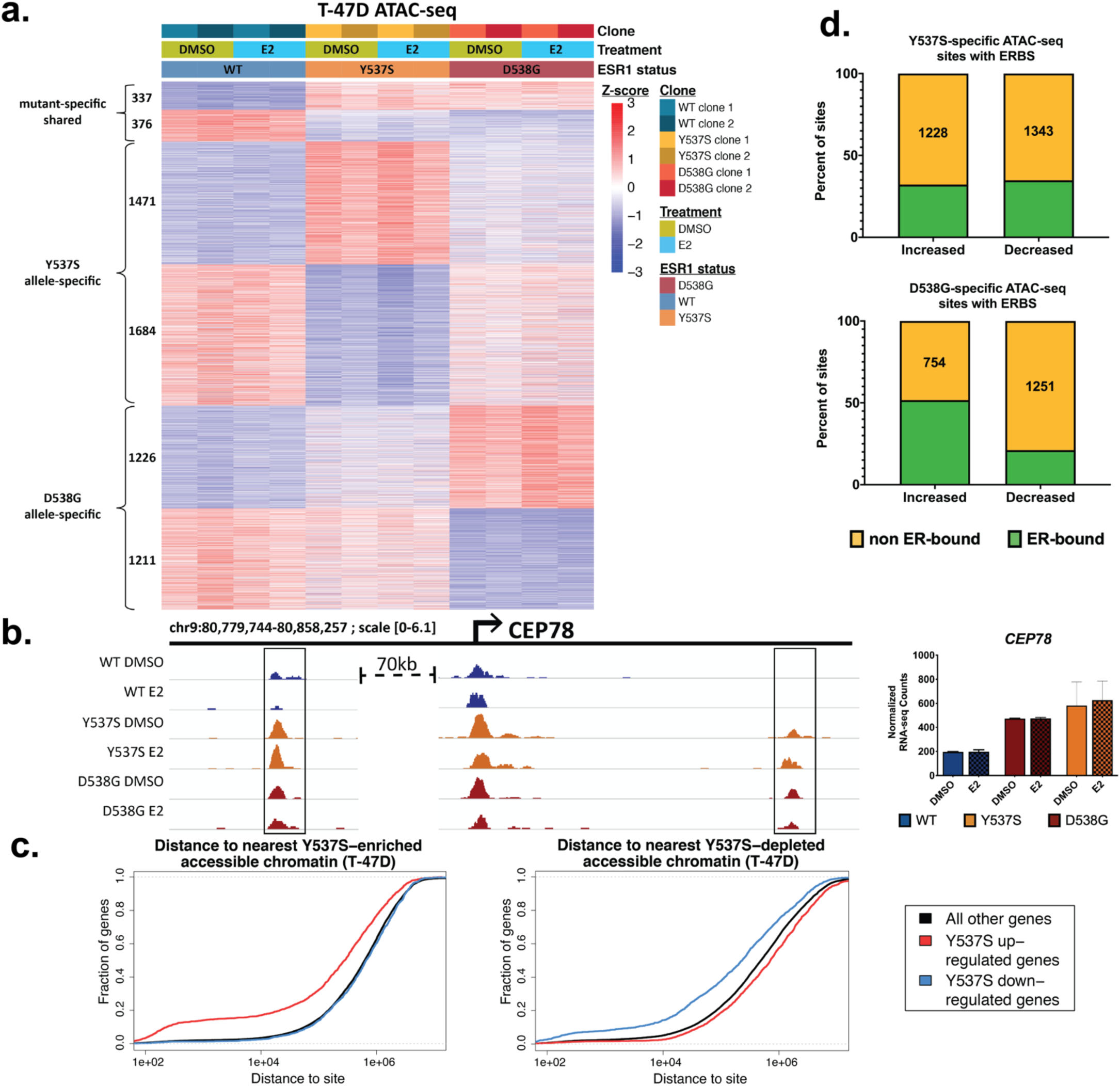
ER mutant cells exhibit large-scale alterations in chromatin accessibility. (a) Heatmap displays ATAC-seq signal of T-47D mutant-enriched and -depleted chromatin accessibility sites. (b) Example locus of shared mutant-enriched accessible chromatin region is shown. Arrow indicates the TSS of *CEP78*. All tracks are normalized to the same scale. *CEP78* gene expression is shown on the right. (c) Cumulative distribution graphs show distance from Y537S mutant-specific genes to Y537S mutant-enriched accessible chromatin. (d) Percent of mutant-enriched or -depleted accessible chromatin regions that also exhibit ER binding is shown.

We explored the relationship between changes in chromatin accessibility and mutant-specific gene expression by comparing ATAC-seq and RNA-seq results from our T-47D mutant clones. We found that mutant-specific up- and down-regulated genes were significantly enriched near regions of mutant altered chromatin accessibility (Fig. 5c, Supplemental Fig. S9a). Of mutant-specific up-regulated genes 17% above background (all other genes) had at least one mutant-enriched region within 100 kb. Of mutant-specific down-regulated genes, more than 15% above background contained at least one nearby mutant-depleted region (Supplemental Table S5). To illustrate the effect that altered chromatin accessibility could have on gene expression, two regions of increased chromatin accessibility near the mutant-specific *CEP78* gene are shown. These regions correlate positively to the increased expression of the *CEP78* gene and may function as regulatory elements controlling its expression (Fig. 5b). We next compared genes that had nearby mutant-altered ERBS with those harboring nearby mutant-altered accessible chromatin regions and found that many Y537S-specific genes harbored both differential ER binding and accessible chromatin sites within 100 kb of their TSS (Supplemental Fig. S9b). However, chromatin accessibility changes were found much more often nearby mutant-specific genes than changes in ER binding, especially in the context of the D538G mutation. These findings suggest that while some mutant-specific gene expression changes may be a direct consequence of altered ER binding, more often these changes appear to be associated with regions of altered chromatin accessibility and potentially other transcription factors binding these regions.

### Transcription factors associated with regions of differential accessibility in ER mutant cells

To identify additional factors associated with mutant-enriched accessible chromatin, we performed motif enrichment analysis on mutant-enriched ATAC-seq sites, identified above, that were negative for ER binding (ERBS-neg). We also excluded any sites that were proximal to gene TSS (within 2 kb) from our analysis due to the heavy over representation of GC rich regions that could bias motif analysis results. Motif analysis of mutant-enriched ATAC-seq sites resulted in a number of motifs that were significantly enriched in one or both of the mutant genotypes (Fig. 6a, Supplemental Fig. S8c). In D538G-enriched sites, the forkhead factor (FOX) motif was most highly enriched (p-value = 1.5e-31). Y537S-enriched sites were highly enriched for POU (OCT) factor (p-value = 8.0e-78) and CTCF (3.3e-50) motifs. Since the FOX and OCT motifs can bind to many related transcription factors, we analyzed expression data and found that 4 FOX factors and 3 OCT factors had detectable mRNA (Fig. 6b); however, FOXA1 exhibited much higher expression than other FOX factors and OCT1 (POU2F1) was the highest expressed OCT factor. OCT1 was also more highly expressed in ER mutant cells compared to WT. These observations led us to examine OCT1, FOXA1, and CTCF in more depth.

**Figure 6.**
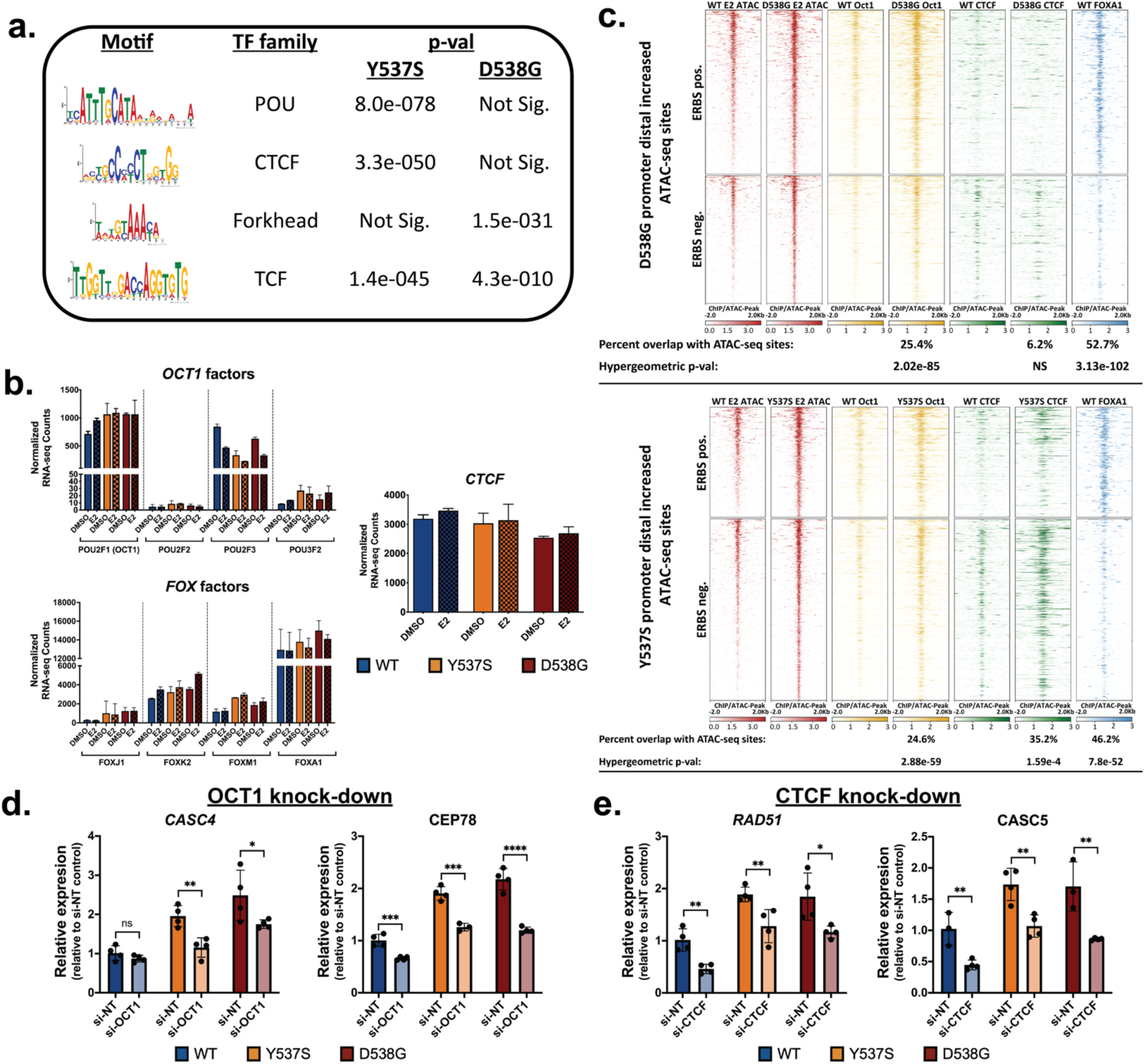
Transcription factors are observed at T-47D mutant-enriched accessible chromatin regions. (a) Enriched transcription factor DNA binding motifs are found at T-47D mutant-enriched accessible chromatin. (b) Gene expression is shown for *CTCF*, *FOX* family genes, and *POU* family genes. Bars represent RNA-seq normalized counts ± SEM for two clones for each genotype and each treatment. (c) Heatmaps display the intensity (depth) of accessible chromatin (red) or DNA-binding of three factors: OCT1 (yellow), CTCF (green), and FOXA1 (blue) at mutant-enriched accessible chromatin regions. qPCR analysis of mutant-specific genes is shown after 72 hours of siRNA-mediated knockdown of *OCT1* (d) or *CTCF* (e). Bars represent average expression ± SEM of two replicates for two clones for each genotype and each treatment. Student’s two sample t-test was used: ^*^p<0.05, ^**^p<0.01, ^***^p<0.001, ^****^p<0.0001, and n.s. = not significant.

To investigate the potential role of these factors in mutant ER cells, we determined the extent to which these factors bind to mutant-enriched accessible chromatin. We performed ChIP-seq for OCT1 and CTCF in the T47-D WT and ER mutant cells in hormone-depleted conditions. We also analyzed FOXA1 ChIP-seq data collected previously in T-47D WT cells (37). OCT1 binding was highly enriched in TSS-distal, mutant-enriched accessible chromatin regions in both the Y537S (p-value = 2.88e-59; hypergeometric test) and D538G (p-value = 2.02e-85 mutant clones. Significant OCT1 binding was found at 25% of both ERBS-negative and ERBS-positive accessible chromatin regions (Fig. 6c). CTCF showed significant enrichment in Y537S mutant-enriched, TSS-distal accessible chromatin (p-value = 1.59e-4) and overlapped with 35% of these sites, but was not enriched in D538G-enriched accessible regions, matching the motif pattern. CTCF was more prevalent at ERBS-negative than at ERBS-positive regions. FOXA1 was enriched in TSS-distal accessible chromatin regions in both Y537S (p-value = 7.8e-52 and D538G (p-value = 3.13e-102 mutant clones and was found at approximately half of both mutants’ TSS-distal accessible regions, but was more evident at ERBS-positive regions than ERBS-negative regions. The enrichment for FOXA1 in ERBS-positive regions is not surprising as it is a known pioneer factor for ER (38) and could be functioning in this role to increase chromatin accessibility in ER mutant breast cancer cells. The binding of these factors at regions of mutant-enriched chromatin accessibility suggests that they could function in altering the expression of mutant-specific genes. Increased CTCF or OCT1 binding in mutant compared to WT cells near mutant-specific genes positively correlates with increased mutant-specific gene expression and suggests a role for these factors in altering gene expression in these cells (Examples in Fig 6d). Overall, the chromatin accessibility analysis has led us to three transcription factors (OCT1, CTCF, and FOXA1) that bind to regions of the genome that become more accessible in ER mutant cells, indicating that these factors may contribute to the unique gene expression patterns attributed to mutant ER.

To determine if OCT1 and CTCF binding to regions of increased accessibility has a functional impact on gene expression, we performed *OCT1* and *CTCF* knockdown experiments. We selected 4 and 6 mutant-specific genes that harbored nearby regions of increased chromatin accessibility and increased CTCF or OCT1 occupancy, respectively. We performed siRNA-mediated knock-downs of *OCT1* and *CTCF*, achieving approximately 50% protein knockdown of CTCF and 70% protein knockdown of OCT1 (Figure S10a). We found that upon *OCT1* knockdown, the expression of 4 of the 6 genes showed a significant reduction in gene expression (Figure 6d and Figure S10b). Two genes exhibited significantly reduced expression in the context of both mutations while two genes had significant reduction in expression in the context of one mutation, although non-significant trends were observed for some genes. With *CTCF* knockdown 2 genes exhibited significantly lower expression in the context of both mutations while the other 2 genes showed no expression effects (Figure 6e and S10c). In the WT context, 1 gene was significantly reduced by *CTCF* knockdown and 2 genes were significantly reduced by *OCT1* depletion. Loss of expression in both the WT and mutant settings indicates that these factors may have a general role in the expression of these genes regardless of ER genotype; however, we found that the effects of *CTCF* and *OCT1* knockdown on expression were larger in ER mutant cells. Overall, these results suggest that CTCF and OCT1 play some role in regulating mutant-specific genes, with OCT1 potentially playing a larger role.

## Discussion

In breast cancer, ER mutations arise under the challenge of hormone therapy that aims to block estrogen production or disrupt ER activity through binding to the LBD. From previous studies it is clear that mutations in ER’s LBD confer ligand independent activity to the receptor(4,5,10,12,14–16), which in large part explains the clinical observation of ER mutations in metastatic breast cancer. However, it has been reported that ER mutations cause additional gene expression changes that appear unrelated to ER’s usual target genes(17–19). In this study, we aimed to confirm ER mutant’s ability to regulate genes that are not normally E2 responsive. By creating isogenic models of the Y537S and D538G mutations in MCF-7 and T-47D breast cancer cell lines, we found that thousands of non-E2 responsive genes changed expression due to the mutation in addition to hundreds of E2 responsive genes that exhibited ligand independent regulation in mutant cells. The genes that changed expression were related to increased growth and migration/invasion, phenotypes that were observed in the ER mutant cells. The increased proliferation may be mediated through E2F transcription factors, based on expression and motif analysis. To determine if mutant ER gene expression effects were specific to our clones or consistently observed across multiple models from different studies, we performed an aggregate gene expression analysis with a total of 16 lines for T-47D and 18 lines for MCF-7 from 3 studies. When taking into account study-to-study variation, we identified thousands of genes that consistently change expression due to ER mutation and do not change expression in response to 8-24 hours of E2 treatment. We observed that most genes have different responses to Y537S and D538G, confirming that allele-specific changes in gene expression are consistent across models from different studies. In addition, we found a significant overlap between the ER mutant effects in cell lines and genes that are differentially expressed between ER mutant and ER wildtype metastatic breast tumors.

One possible explanation for the effect of mutant ER on genes that are not normally E2 responsive is that constant ER activity could have different gene expression consequences than short-term E2 treatments. In an effort to understand the contribution of constant ER activity to the unique ER mutant gene expression patterns, we treated WT cells with E2 for up to 25 days and performed RNA-seq at 5-day intervals. Prolonged exposure to E2 explained approximately half of the mutant-specific gene expression effects with Y537S being more similar to long-term E2 treatment than D538G reiterating the differences unique to each mutant allele. Our findings indicate that approximately half of the ER mutant gene expression effects that appear unrelated to normal ER regulation are in fact estrogen target genes that rely on long-term E2 exposure.

Another way to interpret our prolonged E2 treatment experiment is that half of ER mutant regulated genes cannot be attributed to long-term E2 exposure and must involve novel function of the mutant receptor. We first analyzed genomic binding of mutant and WT ER taking advantage of the FLAG epitope tag that we introduced in the native ER locus. While we observed ligand-independent ER binding in mutant ER cells at the majority of ERBS, we did not observe large-scale reprogramming of ER genomic binding, where approximately 10% of ER bound loci were significantly altered by the mutations. These findings were in contrast to our recent studies on the D538G mutation in endometrial cancer cells that revealed more substantial changes in ER binding (46% of ER bound sites were significantly affected)(22). This indicates that the manner in which mutations influence ER could be unique to different cell types or that ER genomic binding is more robust to mutation in breast cancer cell lines. Both mutant-specific alterations in ER genomic binding(18) and lack of alterations(12) have been reported and our study supports the conclusion that mutations have a relatively small impact on ER genomic binding. While the overall picture of ER genomic binding does not dramatically change with mutation, we still found that approximately 10% of Y537S expression changes could be explained by significantly altered ER binding in ER mutant cells. D538G changes in ER binding were minimal in both T-47D and MCF-7 lines, suggesting that the Y537S mutation may have more of an impact on ER’s genomic binding site selection. Looking at constant ERBS, we found a 10% enrichment near mutant specific genes above background; however, this may be an underestimate of constant ERBS impact on mutant-specific expression due to the large number of constant ERBS, which leads to a high likelihood of a gene having an ERBS nearby simply by chance.

Considering the modest changes in ER genomic binding, we looked for other regulatory regions and factors that could contribute to the unexpected mutant-specific gene expression patterns. By performing ATAC-seq in our WT and ER mutant clones, we found that thousands of loci exhibited differential chromatin accessibility in ER mutant cells. We were able to show consistency in chromatin accessibility changes by analyzing another set of ER mutant clones from a different study (17). Some of these changes in chromatin could be a direct result of mutant ER binding; however, the majority of differentially accessible regions did not harbor ER binding. In addition, differentially accessible regions were at least twice as likely to be found near mutant-specific genes as differential ERBS and could explain approximately 15% of mutant-specific up- and down-regulated genes. These observations led us to the conclusion that other transcription factors contribute to the mutant-specific chromatin landscape and gene expression program.

Motif analysis of differential ATAC-seq sites uncovered OCT, forkhead, and CTCF motifs. Additional experiments confirmed that OCT1, FOXA1, and CTCF were bound at a significant fraction of differentially accessible regions, suggesting that they may be playing a role in increasing chromatin accessibility or possibly taking advantage of new open regions. FOXA1 is a critical factor for ER genomic binding in breast cancer(38) and previous reports have suggested that FOXA1 may not bind to mutant-specific ER bound sites(18). However, our data reveals that FOXA1 could be playing a role in mutant-specific gene expression. OCT1 has been shown to both negatively and positively impact gene expression depending on its associating factors. It can both maintain a poised state of gene expression by facilitating the removal of repressive histone marks and drive a repressed gene expression state by failing to facilitate the removal of these marks(39). In its role as a transcription factor, OCT1 has been shown to contribute to increased cell proliferation, cellular stress response, altered metabolism, regulation of the immune response, and invasion and metastasis(40,41). OCT1 also regulates somatic and cancer stem cell phenotypes in normal and tumor cells(42). OCT1’s presence at mutant-enriched accessible chromatin and increased expression in ER mutant cells suggest that it may be playing a regulatory role and contributing to the expression changes caused by mutant ER. CTCF was found specifically at the Y537S enriched ATAC-seq sites. CTCF plays a key role in the 3D organization of the genome, as reviewed in Ong et al(43), and the presence of CTCF at differential ATAC-seq regions indicates that 3D genome architecture could be altered in ER mutant cells. Depletion of *CTCF* and *OCT1* in ER mutant cells revealed a role for these factors in the regulation of some ER mutant-specific genes. The fact that some mutant-specific genes did not change expression with *OCT1* or *CTCF* knockdown, indicates that many factors may be contributing to the unique gene expression program observed in ER mutant cells. Overall, our analyses of ER genomic binding and chromatin accessibility can explain approximately a quarter of mutant-specific gene expression effects and prolonged ER activity can explain an overlapping 50% of mutant-specific genes. This indicates that additional gene regulation mechanisms are contributing to the ER mutant gene expression program, which may include changes in the 3D genome architecture, post-transcriptional changes, and differences at ERBS, possibly through differential recruitment of cofactors.

Our findings show that the majority of gene expression patterns as well as a small portion of ER genome binding sites are unique to either the Y537S or D538G mutations. Additionally, we see clear allele-specific changes in chromatin accessibility and accompanying transcription factors. Previous studies have also demonstrated allele-specific differences between ER mutations that appear in the structures and molecular effects of mutant ER including differences in their ability to bind cofactors and in their response to hormone therapies(5,6,11,13,17,18,44). These unique differences between ER mutants have significant clinical relevance as *ESR1* mutational status could be used to determine treatment strategies that would best serve the patient. This is particularly true as novel therapies are identified that more effectively treat breast cancers harboring specific *ESR1* mutations. Overall, our study shows that ER mutations consistently impact the expression of thousands of genes that are not normally estrogen regulated and do so partially through constant ER activity and partially through the use of novel regulatory elements.

## Acknowledgments

This work was supported by a DOD Breast Cancer Research Program Breakthrough Award (BC151357) to J.Gertz and J.K.Richer, a DOD Breast Cancer Research Program Expansion Award (BC181341) to J.Gertz, NIH R01CA221303 to S.Oesterreich, and the Huntsman Cancer Institute. Research reported in this publication utilized the High-Throughput Genomics Shared Resource at the University of Utah and was supported by NIH/NCI award P30 CA042014. We thank K-T Varley and Dan Savic as well as Gertz, Varley, Richer, and Oesterreich laboratory members for their helpful comments on the study and the manuscript.

