## Supplemental Methods, Figures, and Tables S2 and S4 - S6 for "Estrogen receptor alpha mutations in breast cancer cells cause gene expression changes through constant activity and secondary effects"

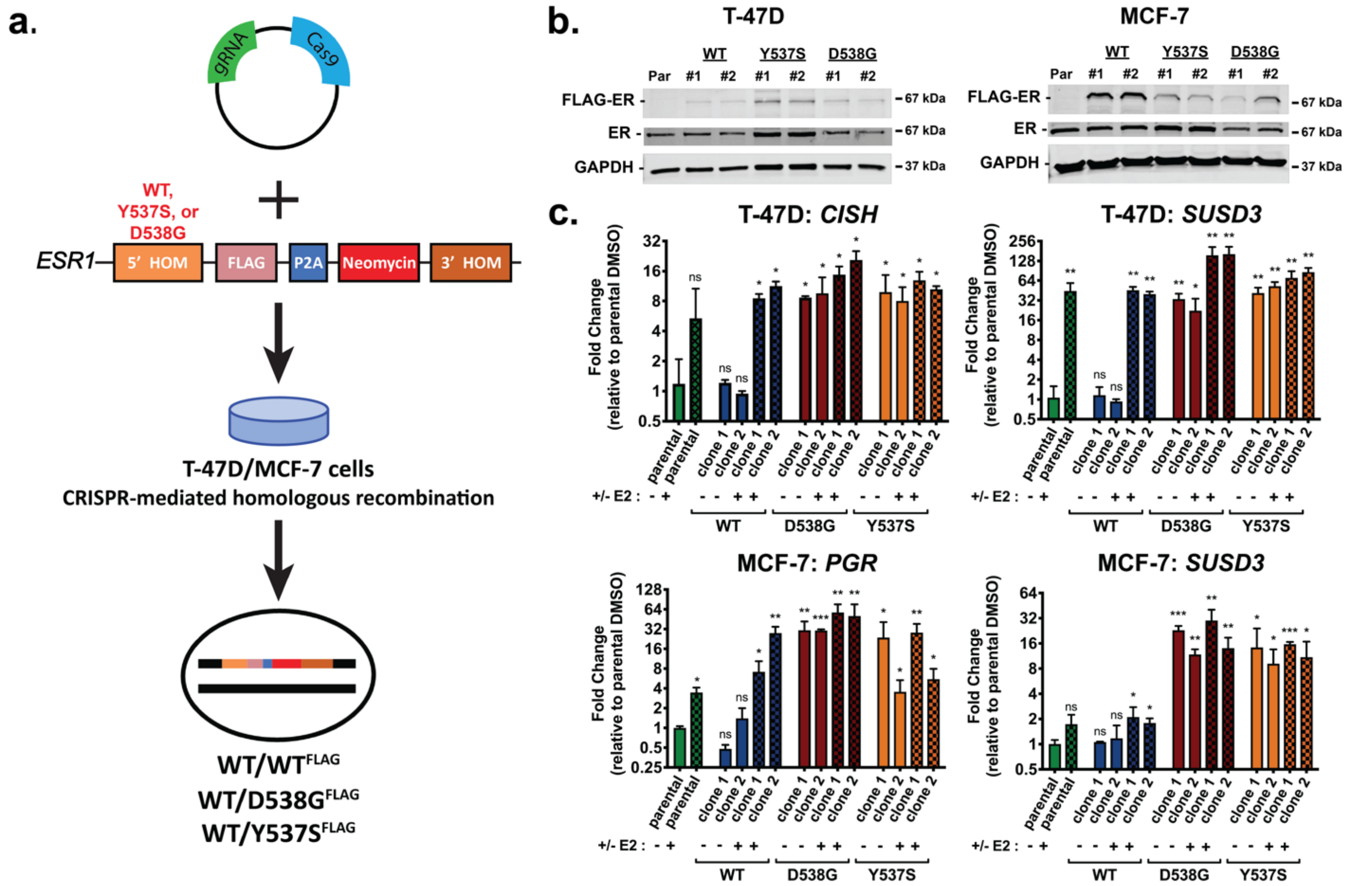


**Supplemental Figure S1. Generation and validation of wildtype and mutant ER breast cancer lines.** (a) Schematic of a CRISPR-mediated epitope tagging strategy that we used to generate multiple heterozygous FLAG-tagged estrogen receptor α (ER) wildtype (WT) or mutant Y537S or D538G clones in T-47D and MCF-7 cell lines. (b) Immunoblots show expression of FLAG, ER and GAPDH, in T-47D and MCF-7 parental, FLAG-tagged WT, Y537S, and D538G ER clones. (c) qPCR analysis of known ER target genes validating ligand-independent gene regulation by D538G and Y537S ER mutants compared to parental and epitope tagged WT clones in T-47D cells and MCF-7 cells. E2 treatments are for 8 hours, bars represent average ± SEM, Student's two sample t-test, *p<0.05, **p<0.01, ***p<0.001, and n.s. = not significant.

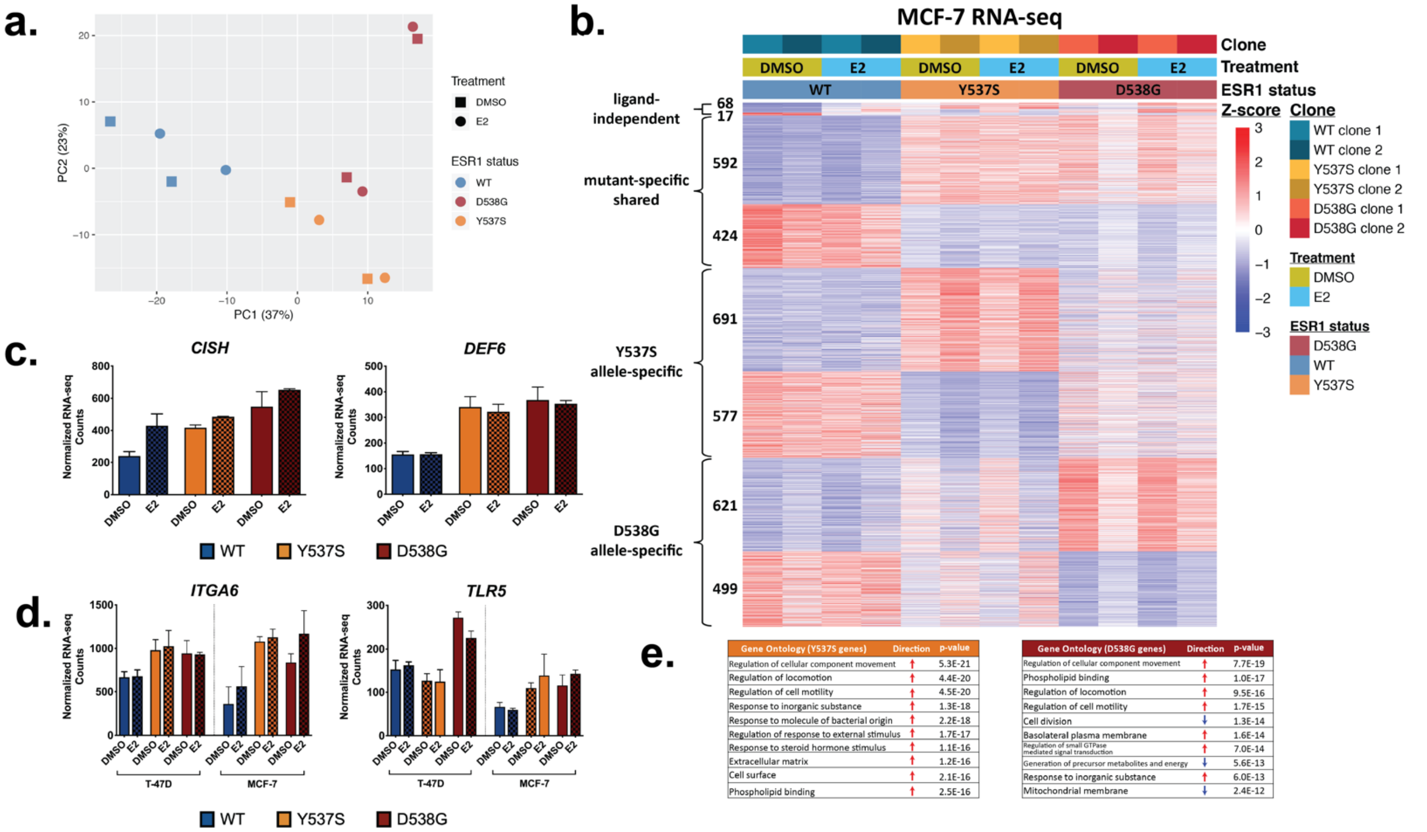


| Gene Ontology (Y537S genes) | Direction | p-value |
|---|---|---|
| Regulation of cellular component movement | ↑ | 5.3E-21 |
| Regulation of locomotion | ↑ | 4.4E-20 |
| Regulation of cell motility | ↑ | 4.5E-20 |
| Response to inorganic substance | ↑ | 1.3E-18 |
| Response to molecule of bacterial origin | ↑ | 2.2E-18 |
| Regulation of response to external stimulus | ↑ | 1.7E-17 |
| Response to steroid hormone stimulus | ↑ | 1.1E-16 |
| Extracellular matrix | ↑ | 1.2E-16 |
| Cell surface | ↑ | 2.1E-16 |
| Phospholipid binding | ↑ | 2.5E-16 |

| Gene Ontology (D538G genes) | Direction | p-value |
|---|---|---|
| Regulation of cellular component movement | ↑ | 7.7E-19 |
| Phospholipid binding | ↑ | 1.0E-17 |
| Regulation of locomotion | ↑ | 9.5E-16 |
| Regulation of cell motility | ↑ | 1.7E-15 |
| Cell division | ↓ | 1.3E-14 |
| Basolateral plasma membrane | ↑ | 1.6E-14 |
| Regulation of small GTPase mediated signal transduction | ↑ | 7.0E-14 |
| Generation of precursor metabolites and energy | ↓ | 5.6E-13 |
| Response to inorganic substance | ↑ | 6.0E-13 |
| Mitochondrial membrane | ↓ | 2.4E-12 |

**Supplemental Figure S2. MCF-7 Y537S and D538G ER mutations exhibit a mutation-specific gene expression profile.** (a) Principal component analysis of MCF-7 RNA-seq data displays the relationships between WT (blue), Y537S (yellow), and D538G (red) MCF-7 clones. (b) Heatmap shows expression of ligand-independent and mutant-specific shared and allele-specific genes identified from MCF-7 RNA-seq analysis. (c) Examples of ligand-independent (*CISH*) and mutant-specific (*DEF6*) gene expression changes are shown. Graphs show average normalized RNA-seq counts. Error bars show standard deviation for two clones for each genotype and each treatment. (d) *ITGA6* and *TLR5* are examples of mutant-specific genes shared across both cell lines (T-47D and MCF-7) for D538G and Y537S mutations, respectively. (e) Gene ontology enriched terms are shown for MCF-7 Y537S- and D538G-specific differentially regulated genes with Fisher's exact test p-values.

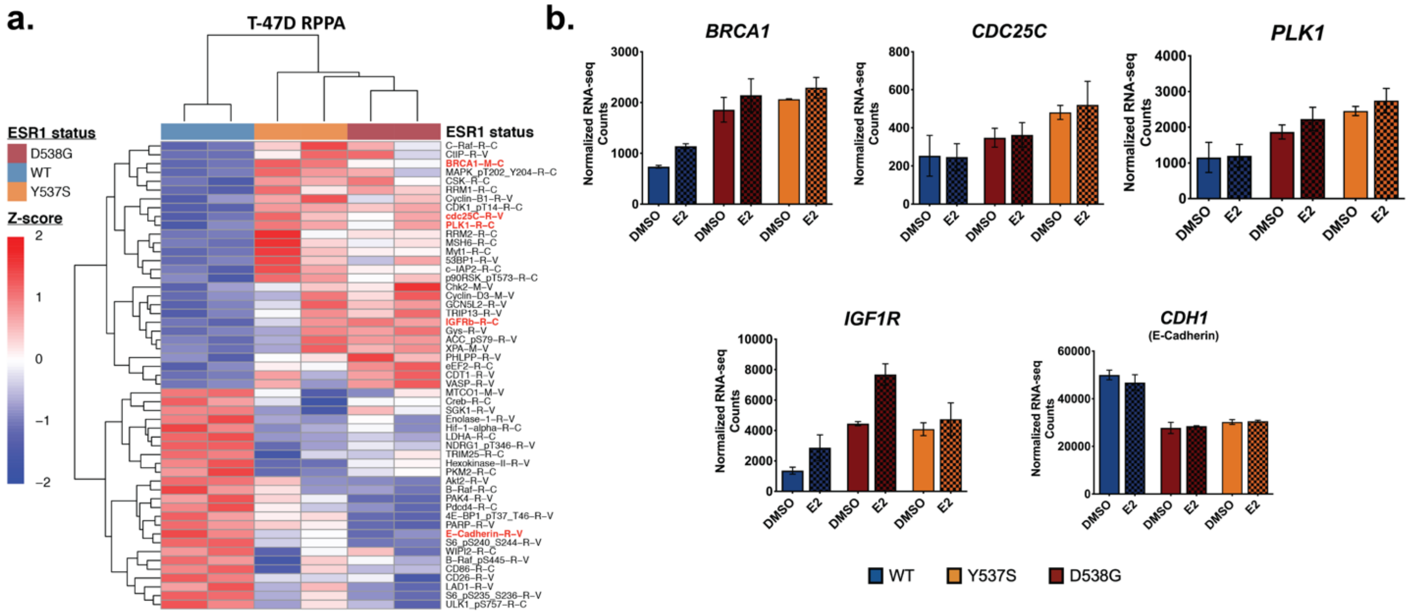


**Supplemental Figure S3. Protein array analysis identifies proteins that are differentially expressed in mutant clones compared to WT clones.** (a) Heatmap shows levels of proteins that are expressed at significantly higher or lower levels in both Y537S and D538G T-47D mutant clones compared to T-47D WT clones. T-test p-values <0.05. (b) Examples of gene expression levels corresponding to protein changes highlighted in (a) are shown. Graphs show average normalized RNA-seq counts. Error bars show standard deviation for two clones for each genotype and each treatment.

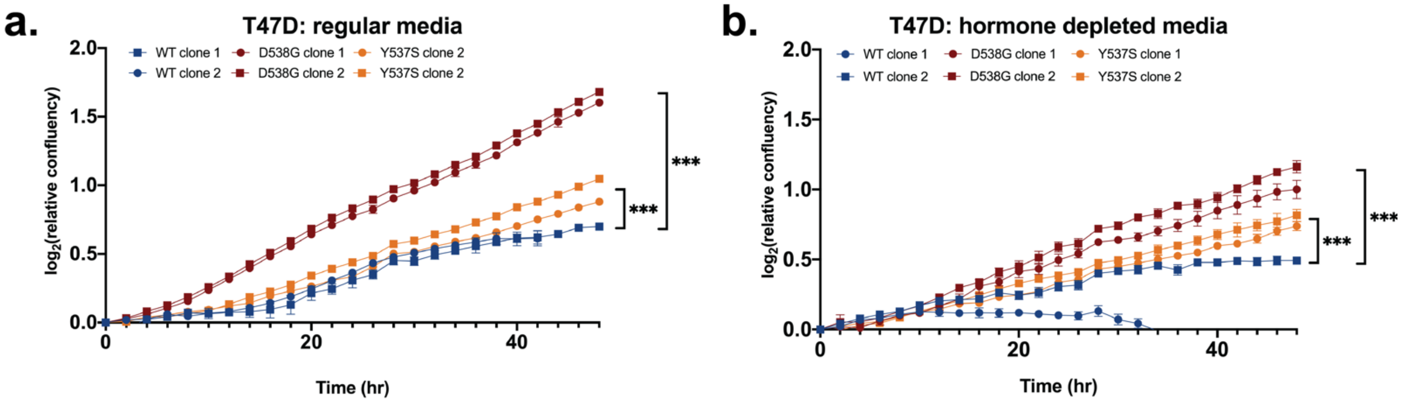


**Supplemental Figure S4. Cell growth analysis of T-47D WT and mutant ER clones.** $Log_2$ of confluency relative to the initial time point is shown for WT and mutant cells in regular media (a) or hormone depleted media (b) over a 48-hour timecourse. Experiments were performed in triplicate. Error bars represent SEM, ***p<0.001.

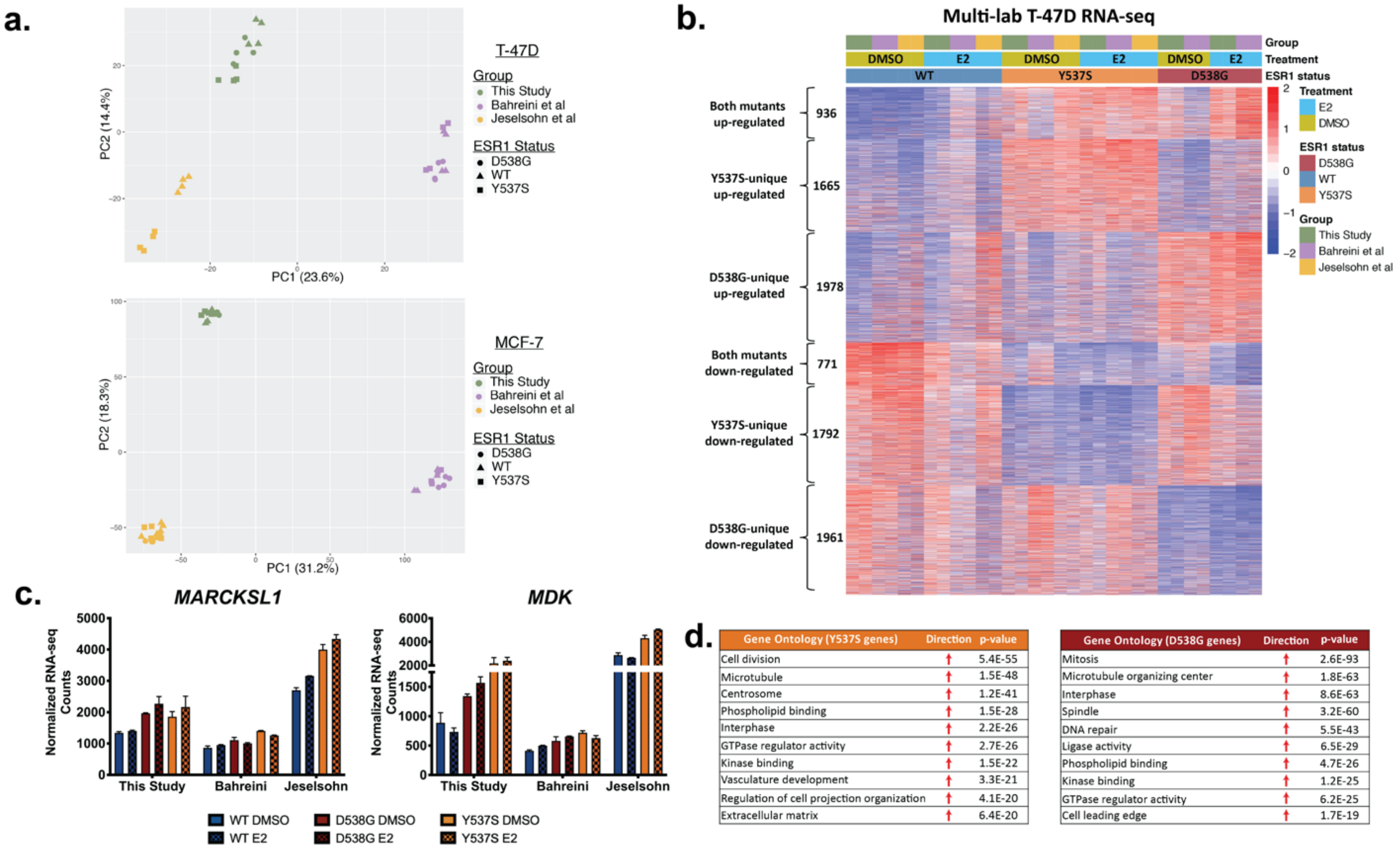


**Supplemental Figure S5. Gene expression analysis of RNA-seq data from three independent groups reveals consistent mutant-specific gene expression patterns in T-47D clones.** (a) Principle component analyses of T-47D (top) and MCF-7 (bottom) mutant and WT clones from three groups shows that the variability between groups exceeds the variability between clones from the same group. (b) Heatmap displays levels of consistent mutant-specific genes. Heatmap columns indicate group of origin, sample type, and treatment for each sample. (c) Examples of genes exhibiting consistent mutant-specific gene expression in T-47D clones are shown. Y-axis shows normalized RNA-seq counts from DESeq2; x-axis groups gene expression by lab of origin and *ESR1* mutational status. (d) Significantly enriched gene ontology terms for Y537S-specific and D538G-specific gene sets from the T-47D multi-lab comparison.

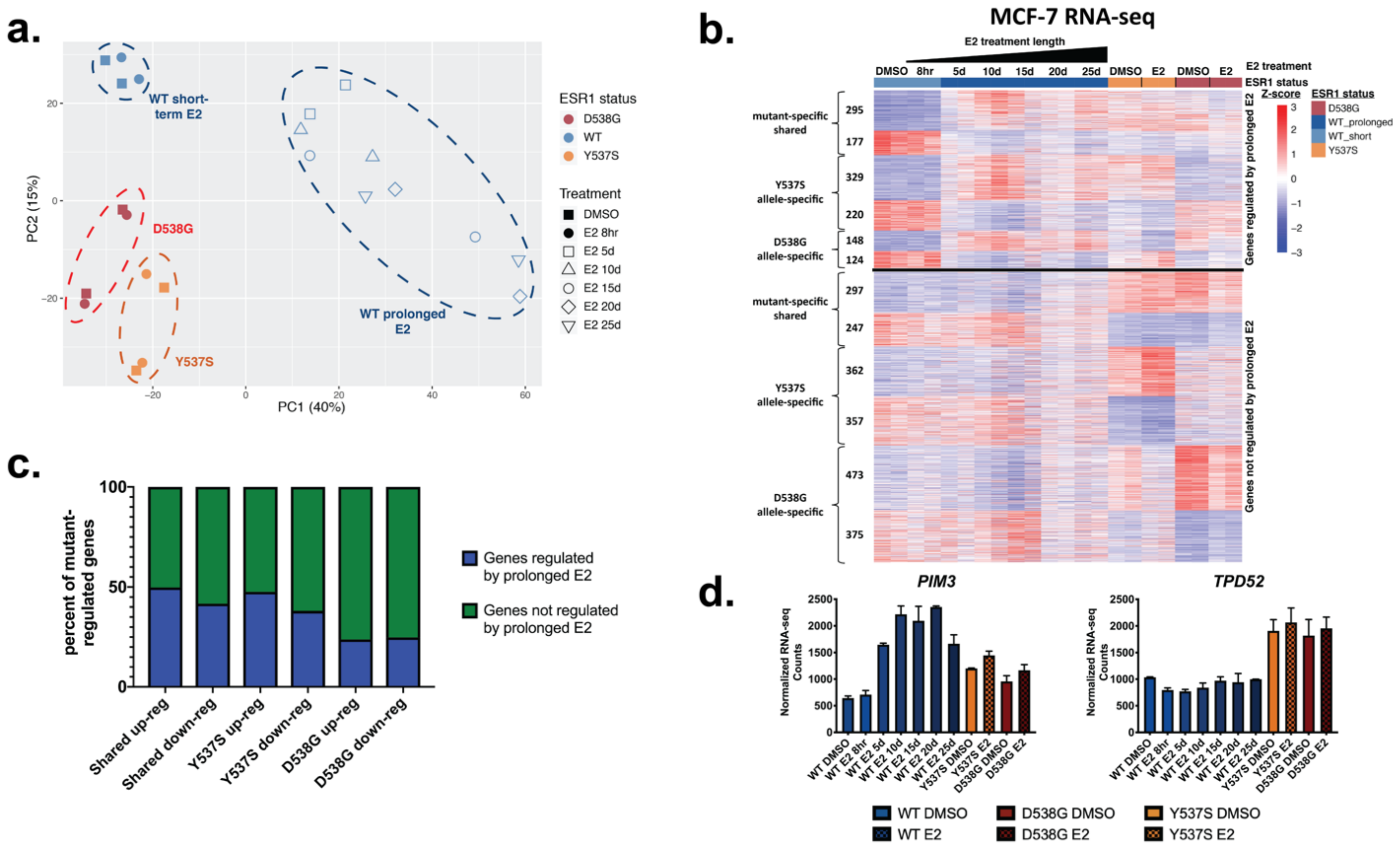


**Supplemental Figure S6. Constant activity of ER explains over one-third of MCF-7 mutant-specific genes.** (a) Principle component analysis of RNA-seq data exhibits the relationships between MCF-7 ER mutant clones and WT clones with short- or long-term E2 treatment. Mutational status is indicated by color and E2 treatment length by shape of symbol. (b) Heatmap shows levels of all MCF-7 mutant-specific genes separated by expression pattern of long-term E2-treatment in WT clones (top: genes regulated by long-term E2 treatment; bottom: genes not regulated by long-term E2 treatment). Mutational status is indicated by color bar and legend. E2 treatment time is indicated above the heatmap. (c) Plot shows the percent of mutant-specific genes regulated by prolonged ER activation in WT clones. (d) Examples of genes that are regulated by long-term E2 treatment (*PIM3*) and that are differentially regulated by mutation only (*TPD52*) are shown.

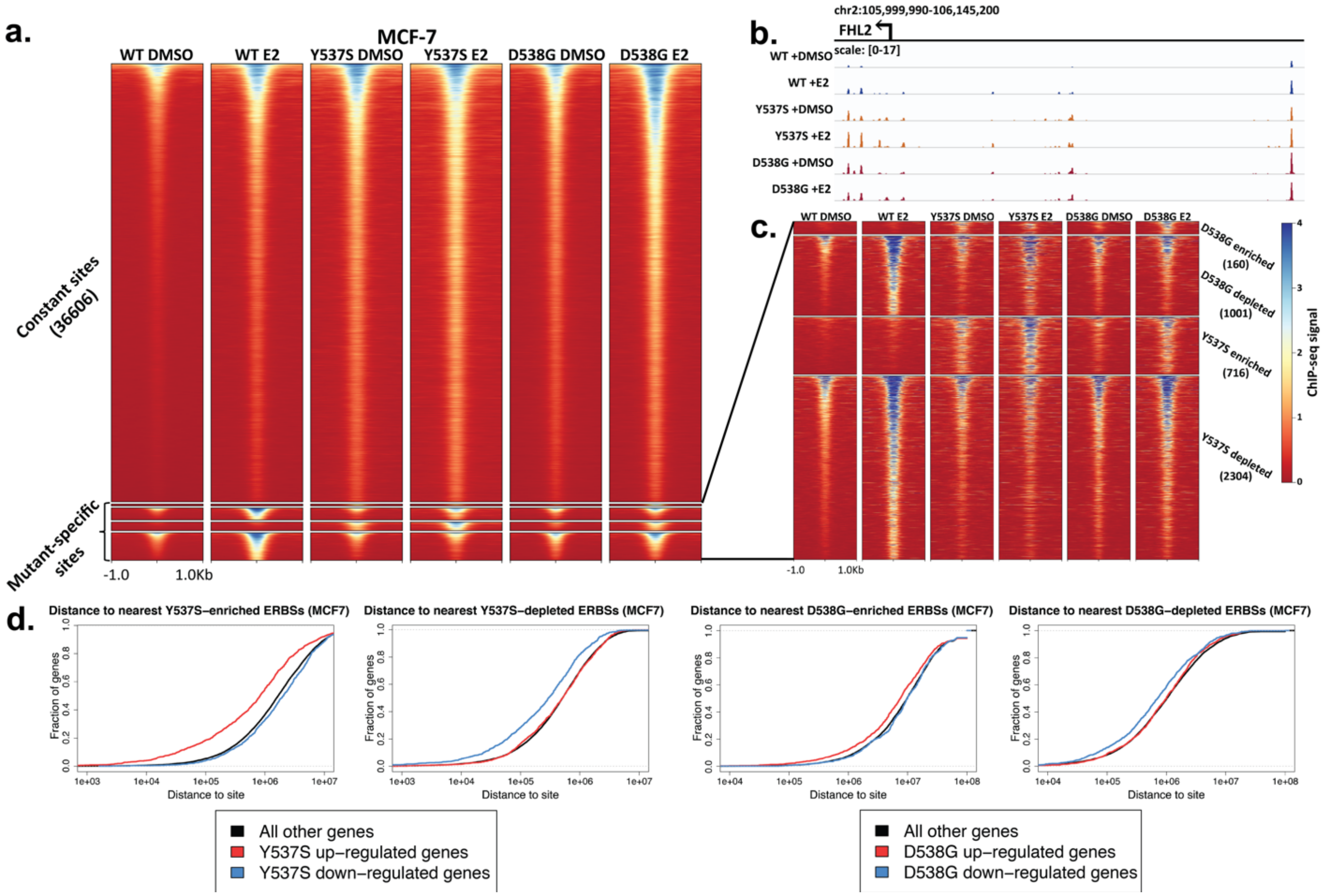


**Supplemental Figure S7. Limited changes in ER's genomic binding are observed in MCF-7 mutants compared to WT and these changes may explain a small portion of mutant-specific gene expression.** (a) Heatmap shows ChIP-seq signal of constant and mutant-specific ERBS in MCF-7 WT and mutant clones with addition of DMSO or E2. (b) Example locus of ligand-independent (constant) ER binding is shown. Arrow indicates the TSS of the *FHL2* gene. All tracks are normalized to the same scale. (c) Heatmap shows ChIP-seq signal of mutant-enriched or depleted sites only. (d) Cumulative distribution plots show the distance from MCF-7 mutant-specific genes to MCF-7 mutant-enriched and -depleted ERBS. Line colors indicate up- or down-regulated mutant-specific genes (red and blue) or all other genes (black).

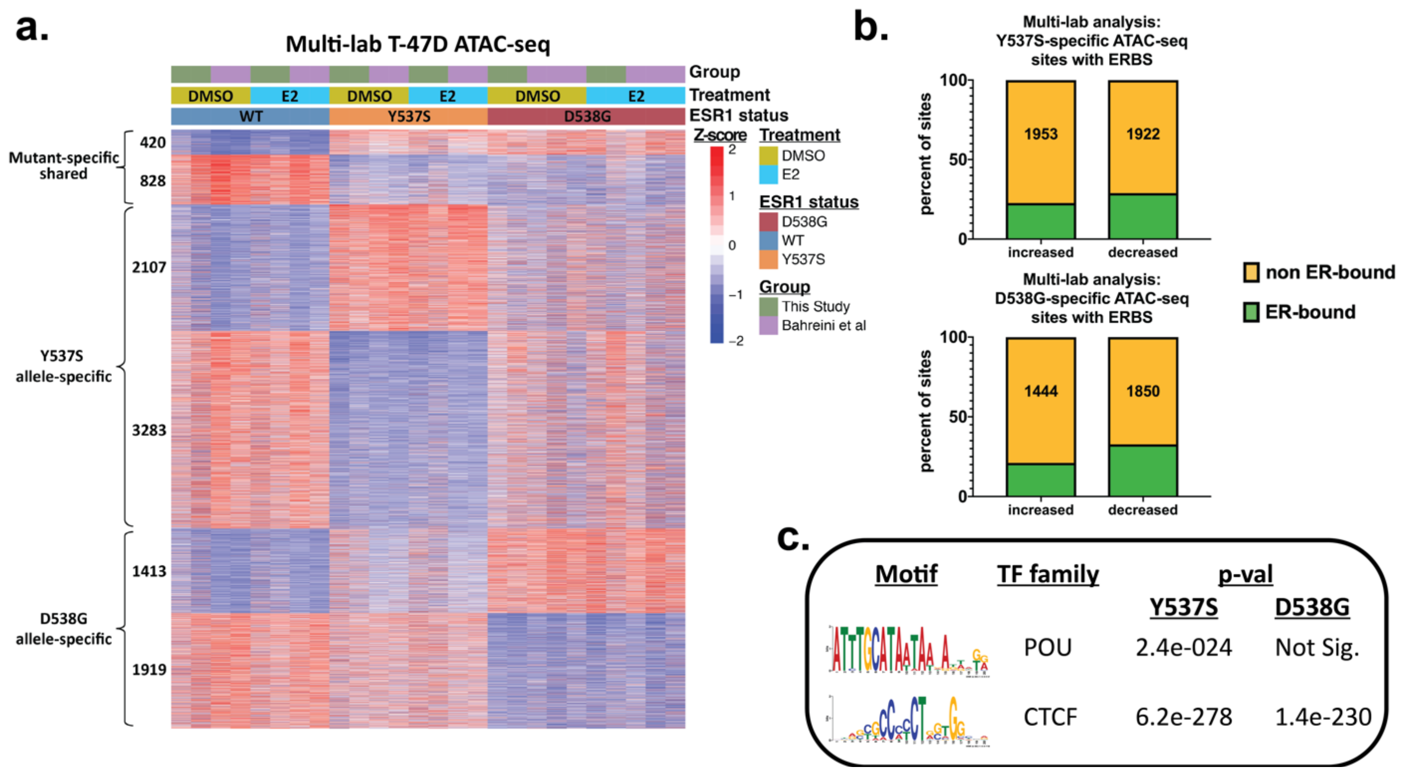


**Supplemental Figure S8. ER mutant cells exhibit large-scale alterations in chromatin accessibility across models from different groups.** (a) Heatmap shows ATAC-seq signal of mutant-enriched and -depleted chromatin accessibility sites from T-47D Y537S and D538G mutant and WT clones from two groups. Differential chromatin accessibility regions are split by those shared across both mutations and those specific to one mutation. Group, treatment, and mutational status are indicated by the header bars and legend. (b) Graphs show the percent of mutant-enriched or -depleted accessible chromatin regions that also exhibit ER binding. (c) Enrichment is shown for transcription factor DNA binding motifs at T-47D mutant-enriched accessible chromatin regions from two groups. Regions queried were TSS-distal and lacked ERBS. Identified motifs are shown (left) along with corresponding adjusted p-values (MEME-Suite E-value) for each mutation.

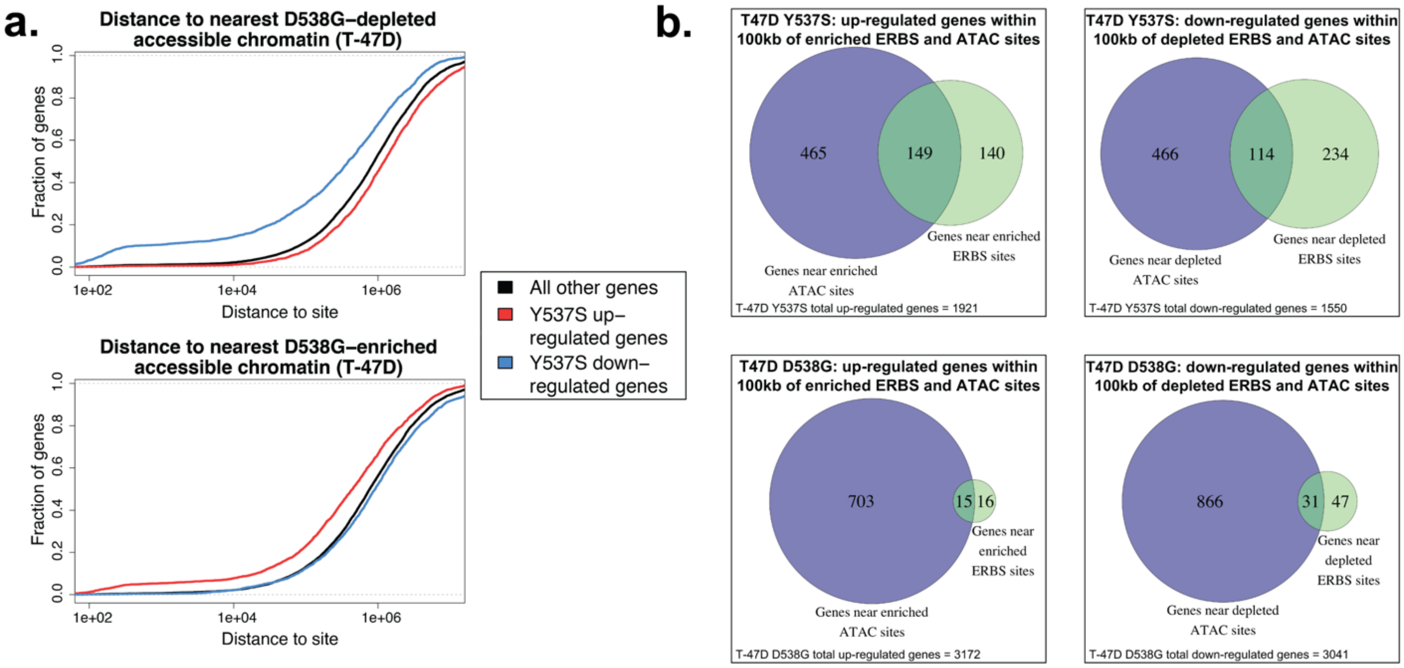


**Supplemental Figure S9. Number of mutant-specific genes that contain either a nearby differential ERBS or ATAC-seq site.** (a) Cumulative distribution graphs show the distance from T-47D D558G mutant-specific genes to D538G mutant-depleted or -enriched accessible chromatin. Line colors indicate up- (red) or down-regulated (blue) mutant-specific genes or all other genes (black). p-values from Wilcoxon tests can be found in supplemental Table S1. (b) Venn diagrams show overlap of genes containing at least one ERBS within 100 kb of their TSS (green) with genes containing at least one increased chromatin accessibility site within 100 kb of their TSS (purple).

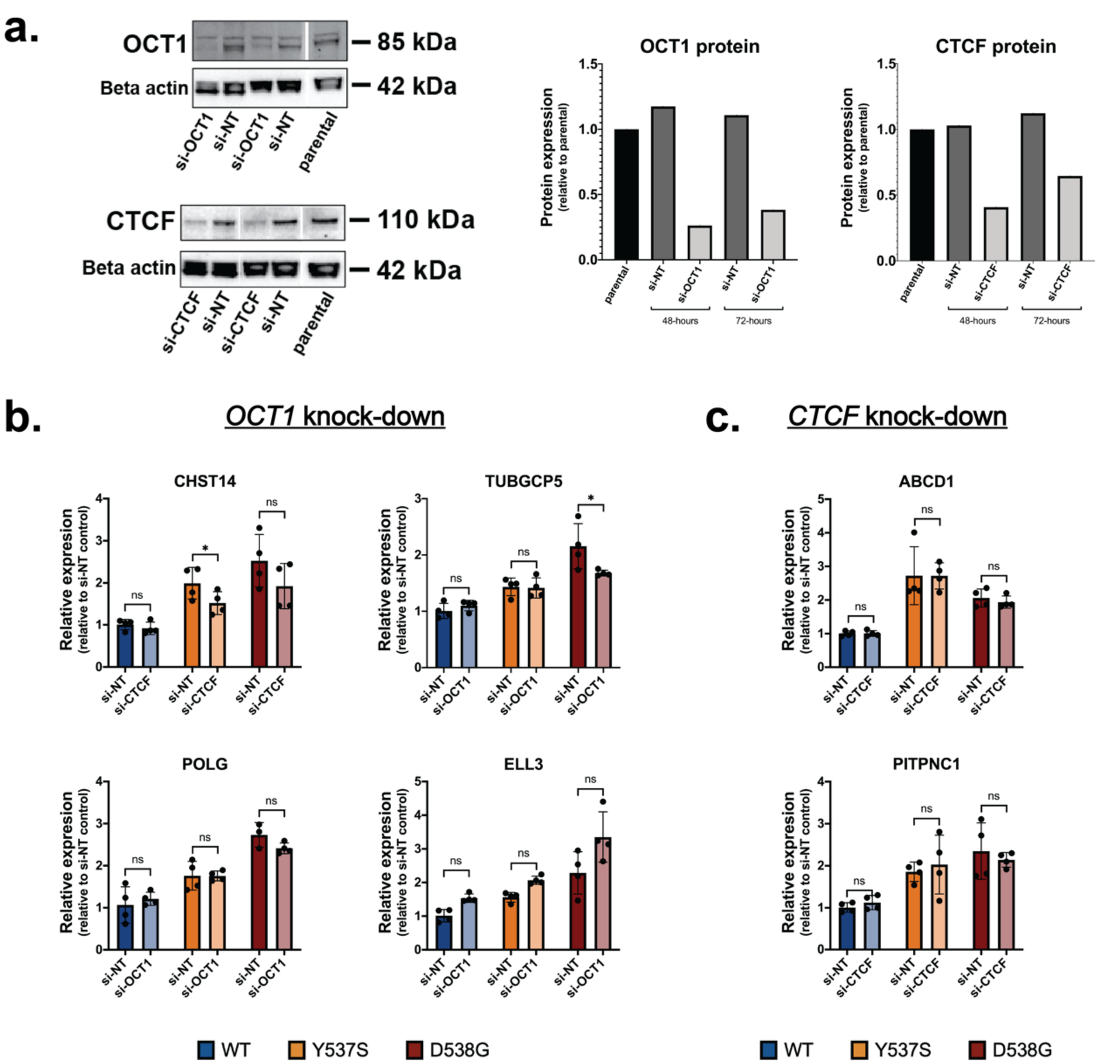


**Supplemental Figure S10. siRNA-mediated knockdown of *OCT1* or *CTCF* effects on mutant-specific gene expression.** (a) Immunoblots show expression of CTCF or OCT1 expression after treatment with either non-targeting (NT) or *CTCF*- or *OCT1*-targeting siRNA (left). Quantification of CTCF or OCT1 protein expression after siRNA treatment (right). (b) Examples of genes with reduced expression in the context of one mutation (top) or neither mutation (bottom) after treatment with *OCT1*-targeting siRNAs. (c) Examples of genes that are unaffected after treatment with *CTCF*-targeting siRNAs. Bar graphs show gene expression relative to WT clones treated with non-targeting siRNA. Bars represent average expression ± SEM of two replicates for two clones for each genotype and each treatment, Student's two sample t-test, *p<0.05 and n.s. = not significant.

**Supplemental Table S2: Comparison of luminal progenitor differential genes with mutant-specific genes.**

| Luminal progenitor gene list | Mutant ER gene list | Number of luminal progenitor genes | Number of mutant ER genes | Number of overlapping genes | % luminal progenitor genes in overlap | P-value |
|---|---|---|---|---|---|---|
| Luminal progenitor up-regulated | T-47D D538G up-regulated | 617 | 3065 | 118 | 19.12 | 0.053445 |
| Luminal progenitor up-regulated | T-47D Y537S up-regulated | 617 | 1859 | 86 | 13.94 | 0.001234 |
| Luminal progenitor up-regulated | MCF-7 D538G up-regulated | 617 | 1083 | 91 | 14.75 | 2.788612e-16 |
| Luminal progenitor up-regulated | MCF-7 Y537S up-regulated | 617 | 1154 | 114 | 18.48 | 3.306366e-26 |
| Luminal progenitor down-regulated | T-47D D538G down-regulated | 714 | 2637 | 95 | 13.31 | 0.799177 |
| Luminal progenitor down-regulated | T-47D Y537S down-regulated | 714 | 1508 | 68 | 9.52 | 0.105766 |
| Luminal progenitor down-regulated | MCF-7 D538G down-regulated | 714 | 807 | 39 | 5.46 | 0.092168 |
| Luminal progenitor down-regulated | MCF-7 Y537S down-regulated | 714 | 875 | 47 | 6.58 | 0.014995 |

**Supplemental Table S4: Overlap between ER mutant associated genes in MET500 study and cell line multi-study analysis.**

| MET500 gene list | Multi-study gene list | Number of MET500 genes | Number of multi-study genes | Number of overlapping genes | Percent of MET500 genes in overlap | p-value |
|---|---|---|---|---|---|---|
| MET500 mutant down-regulated | T47D D538G down-regulated | 160 | 2732 | 18 | 11.25 | 0.393042 |
| MET500 mutant down-regulated | T47D Y537S down-regulated | 160 | 2563 | 21 | 13.13 | 0.096356 |
| MET500 mutant up-regulated | T47D D538G up-regulated | 70 | 2914 | 13 | 18.57 | 0.041589 |
| MET500 mutant up-regulated | T47D Y537S up-regulated | 70 | 2601 | 15 | 21.43 | 0.003007 |
| MET500 mutant down-regulated | MCF7 D538G down-regulated | 160 | 1528 | 18 | 11.25 | 0.005397 |
| MET500 mutant down-regulated | MCF7 Y537S down-regulated | 160 | 2161 | 17 | 10.63 | 0.162928 |
| MET500 mutant up-regulated | MCF7 D538G up-regulated | 70 | 1753 | 14 | 20 | 0.000178 |
| MET500 mutant up-regulated | MCF7 Y537S up-regulated | 70 | 2420 | 11 | 15.71 | 0.053428 |

**Supplemental Table S5. Association between regulatory regions and mutant-specific genes.**

| Differential regulatory regions | % genes harboring a site within 100kb of TSS (p-value) | | | Difference of significant group over background |
|---|---|---|---|---|
| **T-47D Y537S-specific sites** | Y537S up-regulated genes | Y537S down-regulated genes | All other genes | |
| **ER binding enriched** | 15.15% (2.08e-30) | 6.71% (0.008) | 6.90% | 8.25% |
| **ER binding depleted** | 9.81% (0.99) | 22.66% (2.89e-23) | 12.39% | 10.26% |
| **Constant ERBS** | 70.8% (1.06e-14) | 79.9% (9.95e-68) | 64.4% | 6.4% (up)<br>15.5% (down) |
| **ATAC enriched** | 32.25% (2.6e-85) | 13.93% (0.981) | 15.01% | 17.24% |
| **ATAC depleted** | 18.72% (1) | 37.76% (3.06e-34) | 22.5% | 15.28% |
| **T-47D D538G-specific sites** | D538G up-regulated genes | D538G down-regulated | All other genes | |
| **ER binding enriched** | 0.99 (1.75e-32) | 0.27 (1) | 0.67% | 0.32% |
| **ER binding depleted** | 0.95 (1) | 2.60 (8.96e-34) | 1.30% | 1.30% |
| **Constant ERBS** | 55.6% (1.91e-41) | 59.6% (1.59e-71) | 46.6% | 9% (up)<br>13% (down) |
| **ATAC enriched** | 22.85% (1.89e-58) | 12.53% (1) | 13.12% | 9.73% |
| **ATAC depleted** | 7.86% (1) | 28.89% (6.7e-87) | 11.84% | 18.05% |
| **MCF-7 Y537S-specific sites** | Y537S up-regulated genes | Y537S down-regulated | All other genes | |
| **ER binding enriched** | 18.62% (8.81e-52) | 3.96% (1) | 5.42% | 13.20% |
| **ER binding depleted** | 16.64% (0.33) | 28.50% (3.62e-33) | 14.71% | 13.79% |
| **Constant ERBS** | 85.2% (2.32e-78) | 74.8% (2.44e-16) | 66.4% | 18.8% (up)<br>8.4% (down) |
| **MCF-7 D538G-specific sites** | MCF-7 D538G up-regulated | MCF-7 D538G down-regulated | All other genes | |
| **ER binding enriched** | 2.00% (1.27e-9) | 0.66% (0.802) | 0.70% | 1.30% |
| **ER binding depleted** | 9.31% (0.131) | 13.91% (4.22e-11) | 8.50% | 5.41% |
| **Constant ERBS** | 84.9% (2.32e-93) | 77.9% (2.75e-23) | 66.4% | 18.5% (up)<br>11.5% (down) |

**Supplemental Table S6. Oligos used for template amplification to generate Y537S mutant clones, primers for qPCR analyses, and siRNA knockdowns.**

| Name | Sequence |
|---|---|
| gBlock (IDT) with Y537S mutation in red | CTTGAACTGCTTTACTCATTTAAAATACCCA CTCCTGCTTGGCTGAATATCTCATGTTGTCT TTTTAGAAGCTTTGGCGATCCTATTTGAATG CATTTAGGTCCTATTGGAGGGGAATAGGAT CTCATTTGAGGCCACGGAGGTCCATGGAAG TCACCTGCATAGCAAATACCCTGAAAGTGG CTGCAGGGAGAGTGTGAGGGTGGGACCGC CCTGGTAGGAGGTGGAAAATGAAAAACACA CGGCCATGAGTTCCAGATTAGGGCTTCTGA AAGCCCTCAGCTTTCCCAGCTCCCATCCTA AAGTGGGTCTTTAAACAGGAAGAAAGAAAG ATTGCTAAGTGTCTTTGGAGTTCCTCTTCCT TCCCCTTCTAGGGATTTCAGCACTCCTGGG GCTCGGGTTGGCTCTAAAGTAGTCCTTTCT GTGTCTTCCCACCTACAGTAACAAAGGCAT GGAGCATCTGTACAGCATGAAGTGCAAGA ACGTGGTGCCCCTCTCTGACCTGCTGCTG GAGATGCTGGACGCCCACCGCCTACATGC GCCCACTAGCCGTGGAGGGGCATCCGTG GAGGAGACGGACCAAAGCCACTTGGCCA CTGCGGGCTCTACTTCATCGCATTCCTTG CAAAAGTATTACATCACGGGGGAGGCAGA GGGTTTCCCTGCCACGGTCTGA |
| CISH_qPCR_F | TGCCAGAAGGCACGTTCTTAG |
| CISH_qPCR_R | GCCACGAGTGGTTTTCACTG |
| PGR_qPCR_F | ACCCGCCCTATCTCAACTACC |
| PGR_qPCR_R | AGGACACCATAATGACAGCCT |
| SUSD3_qPCR_F | GGTCCCAGCTGAAAGATGAG |
| SUSD3_qPCR_R | GCTCACGGGTTTGTTGAAGT |
| CTCF_qPCR_F | ACCTGTTCCTGTGACTGTACC |
| CTCF_qPCR_R | ATGGGTTCACTTTCCGCAAGG |
| POU2F1_qPCR_F | GTGCATCCAACCACCAATTTG |
| POU2F1_qPCR_R | TGCGTTAGAAGATTTTGCGCTT |
| CASC4_qPCR_F | TCTCTCCCCAAATATGCCTCC |
| CASC4_qPCR_R | AAACGCTGAAGAGGACTGGAA |
| POLG_qPCR_F | CATTCGTGAGAACTTCCAGGAC |
| POLG_qPCR_R | GTGGGGACACCTCTCCAAG |
| ELL3_qPCR_F | CGCTGGTGGTAGCTTGGAC |
| ELL3_qPCR_R | ATGGCTGCCCAAATAATGAGG |
| CEP78_qPCR_F | ACTGGAGGAGTGCCTAAAGCA |
| CEP78_qPCR_R | CAGCTCACTGACTCGTTTATCAA |
| TUBGCP5_qPCR_F | TTCGGGAAACCCTATGGTTACT |
| TUBGCP5_qPCR_R | CAGCACAGATCGTAAACAGCTAT |
| CHST14_qPCR_F | TTTGTGGGCTCCTATGAGAGG |
| CHST14_qPCR_R | GCTGGAAATCGGACGTGAGG |
| RAD51_qPCR_F | CGAGCGTTCAACACAGACCA |
| RAD51_qPCR_R | GTGGCACTGTCTACAATAAGCA |
| ABCD1_qPCR_F | TACCGGGTCAGCAACATGG |
| ABCD1_qPCR_R | TGGTCAGGTTGGAGTAGAGGT |
| PITPNC1_qPCR_F | GATATTGCCTGCGATGAAATTCC |

| | |
|---|---|
| PITPNC1_qPCR_R | AGGCTGATGACTATCTCTCCAG |
| CASC5_qPCR_F | CTTCACACCGAGGACTCAAGA |
| CASC5_qPCR_R | TTTGATGTGTAGAAGAGGCACTG |
| TBP_qPCR_F | CCACTCACAGACTCTCACAAC |
| TBP_qPCR_R | CTGCGGTACAATCCCAGAACT |
| POU2F1 (OCT1)_siRNA_sense | rGrUrCrArArCrArCrCrArArArGrCrGrArArUrUrGrArUrACT |
| POU2F1 (OCT1)_siRNA_anitsense | rArGrUrArUrCrArArUrUrCrGrCrUrUrUrGrGrUrGrUrUrGrArCrUrG |
| CTCF_siRNA_sense | rArArUrUrGrArGrArArCrArUrUrArUrArGrUrUrGrArArGTA |
| CTCF_siRNA_antisense | rUrArCrUrUrCrArArCrUrArUrArArUrGrUrUrCrUrCrArArUrUrGrC |

r = ribonucleotide

### Supplementary methods

#### Quantitative PCR

T-47D and MCF-7 WT or mutant clones were cultured in hormone-depleted media for 5 days and then treated for 8 hours with DMSO or 10nM E2. Cell lysates were harvested using buffer RLT plus (Qiagen) containing 1% beta-mercaptoethanol (Sigma-Aldrich). Total RNA was harvested using a Quick RNA Miniprep Kit (Zymo Research). 50ng of RNA was used per sample to perform qPCR using a Power SYBR Green RNA-to-CT 1-Step Kit (Thermo Fisher Scientific). A CFX Connect Real-Time light cycler (Bio-Rad) was used for thermocycling and fluorophore detection. Expression levels were calculated using the $\Delta\Delta C_t$ method with *CTCF* measurements as the control. Duplicates were measured for each sample. Primers for *CISH*, *SUSD3*, *PGR*, and *CTCF* are found in Supplemental Table S6 Significance of relative expression changes were determined using a one-tailed Student's Two Sample t-test, as implemented in R using t.test, with a p-value cutoff of <0.05.

#### RNA-seq

Cells from T-47D and MCF-7 WT and mutant clones were plated in 100mm dishes and cultured in hormone-depleted media for 5 days followed by treatment with DMSO or 10nM E2 for 8 hours. Cells were washed with phosphate buffered saline (PBS; Thermo Fisher Scientific) and treated with buffer RLT plus (Qiagen) containing 1% beta-mercaptoethanol (Sigma-Aldrich) for cell lysis. Cells were passed through a 21-guage needle and syringe (Sigma-Aldrich) several times to facilitated lysis of genomic DNA. RNA was then extracted using a Quick-RNA Miniprep kit (Zymo Research). RNA was quantified and poly(A) selected RNA-seq libraries were generated using a KAPA Stranded mRNA-Seq kit (KAPA Biosystems) with 500ng RNA per sample as starting material. Libraries were sequenced on the Illumina HiSeq 2500 platform. Resulting RNA-seq data were analyzed to call differential gene expression between WT and mutant clones. Fastq files were aligned to the hg19 human genome build using the HISAT2 spliced aligner (1). SAM files were converted to BAM files using SAMtools (2). Counts for each gene from the University of California Santa Cruz's (UCSC) Known Genes table were assigned using the Subread package's featureCounts program with BAM files as inputs (3). Read counts were normalized and analyzed for differential expression analysis using the DESeq2 package in R (4). RNA-seq was performed for each T-47D and MCF-7 WT, Y537S, and D538G clone. Clones from the same parental cell line and of the same genotype were used as biological replicates in the differential expression analysis between each mutant genotype and the WT clones. PCA plots were generated using the top 5000 most variable genes for each cell line (T-47Ds or MCF-7s). Genes that were significantly up- or down-regulated (adjusted p-value <0.05) in response to E2 treatment (E2-regulated genes) were identified by comparing WT clones treated with E2 to WT clones treated with DMSO. Next, genes that were significantly up- or down-regulated in either the Y537S or D538G mutants were identified by comparing all WT samples to all D538G or all Y537S samples from the same parental cell line regardless of E2 or DMSO treatment. From the resulting differentially expressed genes, E2-regulated genes were removed and the remaining genes were defined as mutant-specific genes. Mutant-specific genes sets were analyzed using Illumina's BaseSpace Correlation Engine to identify pathways and molecular and cellular processes that are significantly associated with mutant-specific gene expression changes.

For the long-term E2 treatment RNA-seq experiments, cells from T-47D and MCF-7 WT clones were cultured in 100mm dishes in hormone-depleted media and were treated with 10nM E2 for 25 days. E2 was added with media changes every 2 days to maintain E2 levels. Cells were taken at 5 day time points (5, 10, 15, 20, and 25 days). RNA purification, RNA-seq library generation, sequencing, and RNA-seq data analysis were all performed as described above. Genes that were significantly differentially expressed in response to long-term estrogen treatment were identified by comparing RNA-seq data from WT clones with DMSO and 8 hour E2 treatments to all WT long-term E2 treatment RNA-seq data. The cutoff for all significant genes was an adjusted p-value of <0.05.

Multi-lab RNA-seq analyses were performed by utilizing publicly-available RNA-seq data from T-47D and MCF-7 WT and Y537S and D538G mutant clones generated by two other groups (5, 6). GEO accession numbers and specific RNA-seq data sets used can be found in Supplemental Table S3. RNA-seq data alignment and count assignment were performed as described above using HISAT2 and featureCounts. PCA plots were generated using the top 10000 most variable genes for each cell line. Differential gene expression analysis was then performed using DESeq2 with a modified multivariate negative binomial regression model to account for variation due to the study in which the RNA-seq data was collected. To implement this multivariate approach, we used the DESeq2 design formula: ~group + E2TreatmentLength + ESR1Status. Y537S and D538G mutant up- and down-regulated genes were identified using an adjusted p-value cutoff of <0.05, which represented significance of the ESR1Status variable.

**Analysis of MET500 data**

RNA-seq fastq files from 91 metastatic breast cancer samples were downloaded from Database of Genotypes and Phenotypes (dbGaP) under accession number phs000673.v2.p1. from the MET500 study (7). Transcript counts from all samples were quantified with Salmon v.0.8.2 (8) and converted to gene-level counts with tximport (9). The gene-level counts from all studies were then normalized together using TMM with edgeR (10). Log2 transformed TMM-normalized counts per million: log2 (TMM-CPM+1) expression values were used for the analysis. To predict ER positivity based on *ESR1* expression, TCGA cohort was used as a reference. Briefly, putative "ER+" (higher than defined cutoff) and "ER-" (lower than defined cutoff) status were predicted based on ESR1 log2 (CPM+1) values of 1045 primary tumors using each consecutive number between 3 (first quartile of ESR1 expression of MET500) and 8.8 (third quartile of ESR1 expression of MET500) with interval of 0.1 individually. The predicted results were then matched to pathological identified ER status under each cutoff selection. Log2 (CPM+1) value of 5.6 was determined as a final cutoff due to its lowest mismatch ratio among all selections (95.5% consistency). 46 putative ER positive samples were then kept MET500 cohort. *ESR1* mutation status was extracted from MET500 portal (https://met500.path.med.umich.edu). 11 ER mutant and 35 ER WT ER+ metastatic tumor samples were used to look for gene that are differentially expressed in tumors with ER mutations. DESeq2 (4) was used to identify differentially expressed genes between the two groups with an adjusted p-value cutoff of 0.05. Genes were required to have a log2 fold change greater than 1.5 or less than -1.5. Significant enrichment of our mutant-specific genes in the MET500 significant gene sets was determined using a hypergeometric test.

**RPPA analysis**

Cells from T-47D WT and mutant clones were plated at 2 million cells per clone on 100mm plates and were cultured in hormone-depleted media for 5 days. Cells were then trypsinized using phenol-red free trypsin and washed twice with PBS. 2 million cells were pelleted and shipped on dry ice for RPPA analysis. RPPA was performed at the MD Anderson Cancer Center Functional Proteomics RPPA Core Facility as described previously (11). In short, protein lysates were serially diluted then spotted onto nitrocellulose-coated slides and probed with 445 antibodies using a tyramide-based signal amplification approach. Slides were stained using a DAB colorimetric reaction and were scanned using a Huron TissueScope scanner. Relative protein levels were determined using a standard curve approach. The resulting normalized log2 protein concentrations were analyzed to identify proteins that exhibited significant increased or decreased expression in either mutant compared to WT. Significance was determined using the Welch t-test, using t.test in R, with a p-value cutoff of <0.05.

**siRNA knock down**

T-47D WT and mutant clones were seeded at 300,000 cells per well in a 6-well format and cultured in hormone-depleted media for 24 hours before transfection with OCT1-targeting, CTCF-targeting, or non-targeting (NT) siRNAs (Integrated DNA technologies) at a 10nM concentration (sequences in Supplemental Table S6). Cells were cultured for 72 hours post transfection and cell lysates were harvested using buffer RLT plus (Qiagen) containing 1% beta-mercaptoethanol (Sigma-Aldrich). Cell lysates were arranged in a 96 deep well plate for total RNA harvesting using a Quick RNA 96 Kit (Zymo Research). Quantitative PCR was performed as described above using primers for *CTCF*, *POU2F1*, *CASC4*, *POLG*, *ELL3*, *CEP78*, *TUBGCP5*, *CHST14*, *RAD51*, *ABCD1*, *PITPNC1*, and *CASC5* (sequences in supplemental Table S6). Expression levels were calculated using the $\Delta\Delta C_t$ method with *TBP* measurements as the control. Experiments were done in duplicate with clones of the same genotype as biological replicates.

**ChIP-seq**

Cells from WT and mutant clones were plated in 150mm dishes and cultured in hormone-depleted media for 5 days. Cells were then treated with either DMSO or 10nM E2 for 1 hour. Following DMSO or E2 treatments, cells were fixed for 10 minutes at room temperature with 1% formaldehyde (final concentration) added directly to the media (Sigma-Aldrich). Fixation was ended by briefly treating media with glycine to a final concentration of 125mM to quench the remaining formaldehyde. Media was removed, and fixed cells were washed once in cold PBS. Fixed chromatin was then harvested by scraping in cold Farnham lysis buffer (5 mM PIPES pH 8.0; 85 mM KCl; 0.5% NP-40) supplemented with protease inhibitor (Thermo Fisher Scientific). Chromatin immunoprecipitation was performed on fixed chromatin as described previously (12) using antibodies against the FLAG epitope tag (Anti-FLAG M2 antibody; Sigma-Aldrich F1804), CTCF (sc-5916, Santa Cruz Biotechnology) or two antibodies against OCT1 combined at equal ratios (A301-716A and A301-717A, Bethyl Laboratories Inc.). Sequencing was performed using the Illumina HiSeq 2500 platform. FOXA1 binding was analyzed using previously obtained FOXA1 ChIP-seq data (13). Resulting reads in Fastq format were aligned to the hg19 human genome build using bowtie2 (14) with the following parameters: --chunkmbs 512 -m 1 -t –best -q -S -l 32 -e 80 -n 2 -p. Output SAM files were sorted and converted to BAM files using SAMtools (2). MACS2 (15) was used to call peaks using a p-value cutoff of 1 x $10^{-10}$ and a mfold parameter between 15 and 100. Control input libraries from WT and mutant lines were used as controls for MACS2 peak calling in each ChIP-seq experiment. Clones were used as replicates for each experiment. Significantly differentially bound FLAG tagged ERBS sites were determined using DESeq2 (4) to compare counts per million at each ER binding site called. FLAG tag ER ChIP-seq results for E2-treated WT clones were compared to either Y537S or D538G DMSO- and E2-treated FLAG tag ER ChIP-seq results. An adjusted p-value cutoff of <0.05 was used to identify significantly mutant-enriched and -depleted ER-bound sites. Constant ER-bound sites were sites that were bound in at least one replicate from any genotype that were not differentially bound according to our DESeq2 analysis. Enrichment of CTCF, OCT1, and FOXA1 binding at mutant-enriched accessible chromatin was assessed using the BEDTools (16) and deepTools (17) packages.

#### ATAC-seq

Cells from T-47D WT and mutant clones, including clones derived in this study and clones reported in Bahreini et al. (5), were plated in 6-well plates and cultured in estrogen-depleted media for 5 days. Following estrogen deprivation, cells were treated with DMSO vehicle control or 10nM E2 for 1 hour, then washed with PBS and removed from the plates using phenol-red free trypsin. Cells were pelleted and resuspended in hormone-depleted media. 250,000 cells for each sample were used for each ATAC-seq experiment. ATAC-seq libraries were prepared as previously described (18) and were sequenced on an Illumina HiSeq 2500 sequencer. Sequenced reads were aligned to the hg19 human genome build using bowtie2 (14) with the following parameters: --chunkmbs 512 -m 1 -t --best -q -S -l 32 -e 80 -n 2. Resulting SAM files were converted to BAM files using SAMtools (2). Regions of accessible chromatin were identified by calling ATAC-seq peaks using MACS2 with an adjusted p-value cutoff of <0.05 and a mfold parameter between 15 and 100. Subread's featureCounts (3) was used to quantify reads that align to all peaks called in any sample. Aligned reads were normalized and mutant-enriched and -depleted chromatin accessibility regions were identified using the DESeq2 package in R (4). Clones developed by our lab were used to identify regions of differentially accessible chromatin. Clones with the same ER genotype were grouped as biological replicates, and all WT samples were compared to all Y537S or all D538G mutant samples to identify mutant-enriched and -depleted chromatin accessibility regions. Overlap of ATAC-seq sites and ERBS identified through FLAG epitope tag ChIP-seq was performed using the BEDTools intersect tool (16) with an overlap size of at least 1 base pair. 100-bp regions surrounding the summits of mutant-enriched non-ER bound ATAC-seq regions >2kb from gene TSS were used to perform motif analysis using MEME-Suite's MEME-ChIP tool (19). Enriched motifs between 6 and 20 bps in length were reported, with a motif site distribution of 0 to 1 occurrence per sequence.

Multi-lab ATAC-seq was performed using ATAC-seq data from T-47D WT and mutant clones developed in this study along with those reported in Bahreini et al. (5). Differential chromatin accessibility was assessed using DESeq2 with a modified multivariate negative binomial regression model to account for variation due to the study in which the clones were developed. A multivariate negative binomial regression model was implemented by using the DESeq2 design formula: ~group + ESR1Status. Y537S and D538G mutant enriched- and depleted-accessible chromatin regions were identified using an adjusted p-value cutoff of <0.05, which represented significance of the ESR1Status variable.

2. Li H, Handsaker B, Wysoker A, Fennell T, Ruan J, Homer N, Marth G, Abecasis G, Durbin R, Genome Project Data Processing S. The Sequence Alignment/Map format and SAMtools. Bioinformatics.

2009;25(16):2078-9. Epub 2009/06/10. doi: 10.1093/bioinformatics/btp352. PubMed PMID: 19505943; PMCID: PMC2723002.
